# Accurate neural control of a hand prosthesis by posture-related activity in the primate grasping circuit

**DOI:** 10.1101/2023.06.02.543242

**Authors:** Andres Agudelo-Toro, Jonathan A. Michaels, Wei-An Sheng, Hansjörg Scherberger

## Abstract

Brain-computer interfaces (BCIs) have the potential to restore hand movement for people with paralysis, but current devices still lack the fine control required to interact with objects of daily living. Following understanding of cortical activity during arm reaches, hand BCI studies have focused on velocity control. However, mounting evidence suggests that posture, and not velocity, dominates in hand-related areas during natural movement. To explore whether this signal can causally control a prosthesis, we developed a novel BCI training paradigm centered on the reproduction of hand posture transitions. Macaque monkeys trained with the protocol were able to control a multi-dimensional hand prosthesis at high-accuracy, including execution of the very intricate precision grip. Subsequent analysis revealed that the posture signal in the target grasping areas was a major contributor to control. Population activity exhibited pattern separation and dimensionality increases driven by posture kinematics, and simulations with a grasping circuit model demonstrated the generalizability of our approach. We present for the first time neural posture control of a multi-dimensional hand prosthesis, opening the door for future devices to leverage this additional information channel.

## Introduction

The primate hand is a fascinating and highly capable organ. From grooming to tying a knot, our hands can rapidly switch among the abundant set of poses required for the prehensile and non-prehensile interaction with objects^1,2^. Disruption of hand function caused by spinal cord injury severely limits this capacity, making it a top recovery priority for patients^3^. By establishing a direct connection to cortex, brain-computer interfaces (BCIs)^4–13^ can potentially drive a large number of degrees of freedom and thus, a fully articulated hand prosthesis. Although significant progress has been made toward this goal^8,14–18^, and high-performance recording techniques and robotics are increasingly available^19–21^, BCI devices are still limited to basic grasping and cannot yet reproduce the rich set of configurations of the native hand.

Despite evidence of abundant posture information in hand-related cortical areas during native grasping^14,16,22–26^, hand BCI protocols have focused on the movement velocity aspect of control, using this approach to control hand opening^6–9^, hand joint synergies^17^, or individual finger movements^15^. Focus on movement velocity^12,13^ follows the traditional view of motor cortex representing movement direction^27–29^ and indications that, at least for BCI reaching, using velocity control yields higher performance^30–32^.

During posture control^11,33,34^, kinematic trajectories in the workspace determine effector state transitions (Fig. 1A,B). Although multiple studies have shown correlational evidence that such kinematic trajectories are encoded during native grasping^16,23,14,35,24,25,22,36,26^, the use of this code has not been demonstrated in a hand prosthesis. Leveraging this posture signal^14,16,23–25^ holds promise for improving future hand neuro-prosthesis by introducing a control channel complementary to velocity^15^.

**Figure 1.**
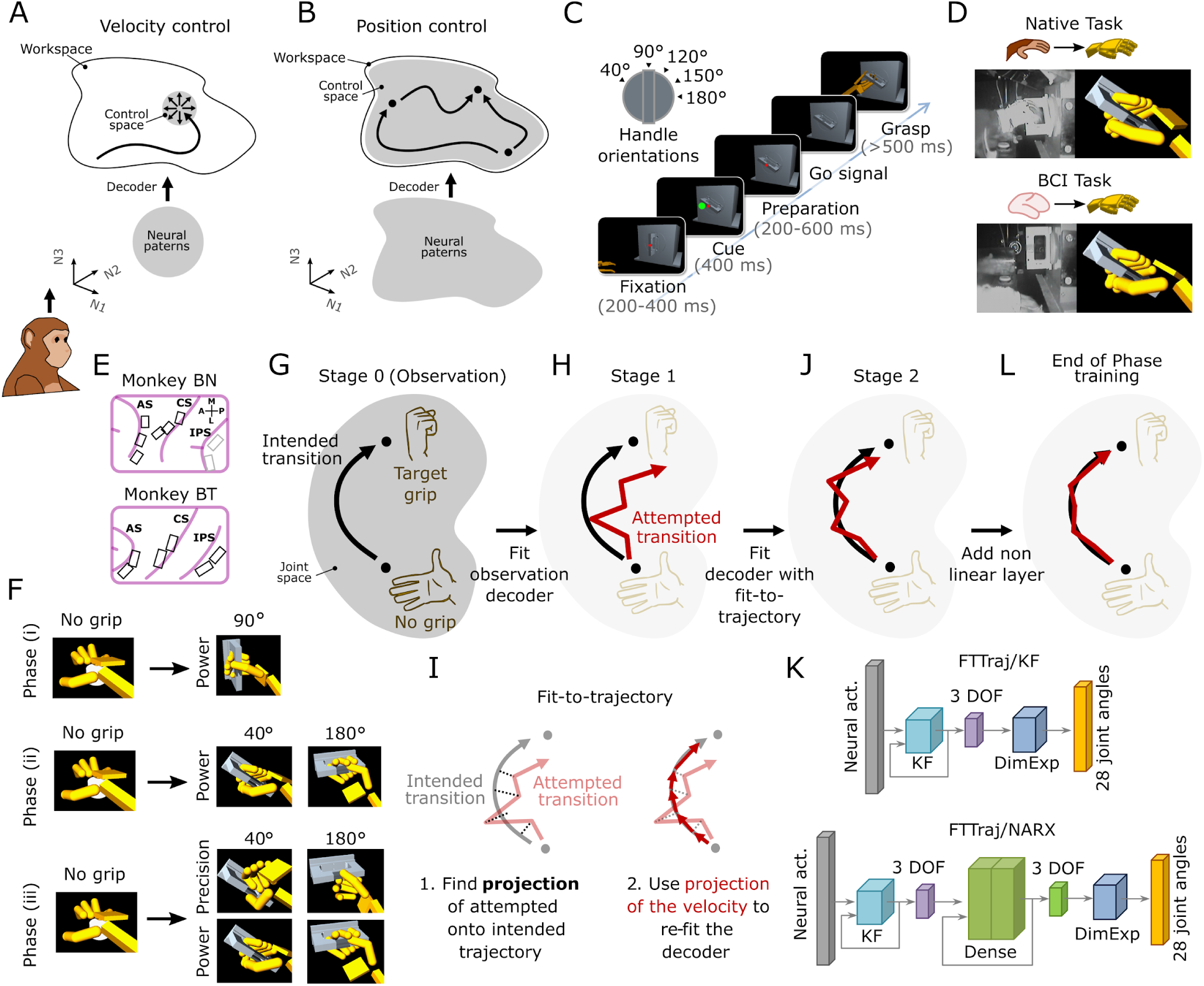
Description of the task and training strategy. (A-B) Velocity vs. position BCI control using a linear decoder. (A) During velocity control (e.g. cursor control), the user controls at each timestep a direction to move next in the workspace (top, circle with arrows). Neural population activity should reflect the different orientations and circle size to cover all possible directions and speeds (bottom). (B) During position control, the user determines at each timestep positions in the workspace. These can be seen as trajectories and states in a space of configurations (top). Population activity (bottom, dashed area) should resemble this workspace. (C) Handle orientations and task flow. The task started with a fixation dot, followed by a cue indicating grip type and handle orientation. After a preparation period, a “Go” signal was indicated by absence of the fixation dot. The animal should then grasp the handle with the virtual hand and hold it for at least 500ms. Monkeys were then rewarded and returned their hand to the initial resting position. (D) Task configurations. Native task: the subject controlled the virtual hand using the native hand. Hand tracking was used to determine the exact movement of the virtual hand. BCI task: the virtual hand was controlled through brain activity while the subject remained still. (E) Neural data was extracted from floating microelectrodes on the hand area of M1, area F5 of the ventral premotor cortex, and the anterior intraparietal sulcus (AIP, only in monkey BT). (F) BCI training phases. In Phase (i), we trained the power grip for the 90° handle orientation. In Phase (ii), the power grip at 40 and 180°. In Phase (iii), both power and precision grips for 40 and 180° (G-J) Daily decoder calibration. (G) At Stage 0, the subject observes fully automated grips while neural activity is captured. Neural activity and the idealized (intended) transitions in joint kinematics space are used to fit an observation decoder. (H) The observation decoder executes a batch of grip attempts (typically with relatively low performance) (I) Apply fit-to-trajectory (FTTraj). The attempted transitions of Stage 1 are projected onto the intended trajectories to obtain the velocity vectors used to fit the decoder. During grip hold periods, zero velocity and the intended grip configuration are assumed. Neural and projected data are used to fit a new decoder. (J) The Kalman filter (KF) decoder fit with Stage 1 data increases the performance of the transitions but it is usually limited in accuracy. (K) A non-linear layer (two layers densely connected NARX) was added after the KF to increase accuracy. Output of the KF or NARX layer was converted from 3D to the 28D joint angle space through a dimensionality expansion step (DimExp, Methods). (L) The final FTTraj/NARX decoder increases accuracy during transitions and hold periods.

In this work, we demonstrate that the cortical posture signal in hand-related areas can support accurate hand neuroprosthetic control for multiple degrees of freedom. To show this, we developed a BCI intention estimation protocol that incorporates position and velocity control through re-fitting to kinematic trajectories. The protocol assumes the subject aims to execute a transition in the space of hand configurations, extending intention estimation for velocity control^37^ to posture-based control.

When tested in two rhesus monkeys implanted in three key areas of the grasping circuit^38^ (AIP, F5, and hand M1), our approach generated accurate hand state-transitions and hold postures for two different grip types in real-time. Post-hoc analyses showed that most of the information required for BCI control was position-like, mirroring native hand use^14,16,23–25^. To support hand state-transitions, the neural population exhibited separation by grasp type and dimensionality increases over training, further supporting the view that cortex encoded the desired posture-based trajectories.

## Results

### Training paradigm

To study cortical control of hand prostheses, we employed a grasping task with visual feedback through a virtual environment (Fig. 1C, Fig. S1A). In the task, subjects were presented with a rotating handle in five possible orientations and asked to grasp the handle using two principal grips^2^ (power or precision). Depending on the task setting, subjects grasped the handle using a hand avatar driven by the native hand (through a tracking device), or the BCI (Fig. 1D). In the BCI setting, subjects executed the task without concurrent grasp movement (lifting the hand from a rest button aborted the trial, Methods). To drive the BCI, subjects were implanted in the anterior intraparietal area (AIP), ventral premotor area F5, and the hand area of primary motor cortex (M1) (Fig. 1E), using floating microelectrode arrays (Microprobe, USA).

Our BCI training approach is detailed in Methods. In short, monkeys were incrementally trained^7,17,39^ to drive the hand avatar through a BCI decoder in three Phases: control of grip aperture (i), control of wrist rotation (ii), and continuous grip selection (iii) (Fig. 1F, Fig. S1, Fig. S2). To fit the decoder, we used a three-stage procedure following closed-loop recalibration and intention estimation protocols (Fig. 1G-J, Methods)^37,40,41^. Contrasting previous target-based recalibration approaches^37^, we re-fitted position-velocity Kalman filter (KF) decoders using a *fit-to-trajectory* approach. This approach centers around transitions instead of targets, attempting to preserve position and velocity features of joint movements (Fig. 1I, Fig. S1, Fig. S2, Methods). This implies all the position points and velocity vectors along the trajectory are used to recalibrate the decoder. By comparison, target based recalibration approaches (*fit-to-target*) only re-fit velocity vectors^37^.

After training our subjects to use the prosthesis with *fit-to-trajectory* (FTTraj/KF), we further boosted grasping accuracy by augmenting the position-velocity KF with a non-linear autoregressive with exogenous inputs (NARX) network^42^ which we applied at the end of each training phase (FTTraj/NARX, Fig. 1K,L).

### Performance overview

We tested our BCI training strategy in two rhesus monkeys (monkey BN and BT) previously trained to perform the grasping task with the native hand. After completing Phase (iii) of the training (Fig. S3A), monkeys were able to fully drive the hand prosthesis achieving the different grip types and handle orientations with high accuracy (Fig. 2A). To assess the accuracy of the BCI grips, we compared the executed virtual grips to native hand grips. Grip accuracy was comparable to native grips when holding the handle (Fig. 2B), as measured by the average single trial position variance during the 500 ms hold period. Transitions to hold pose (i.e. the displacement from open to closed hand in the workspace) were also comparable to native grips when measuring excess trajectory and error to average transitions (Fig. 2C,D).

**Figure 2.**
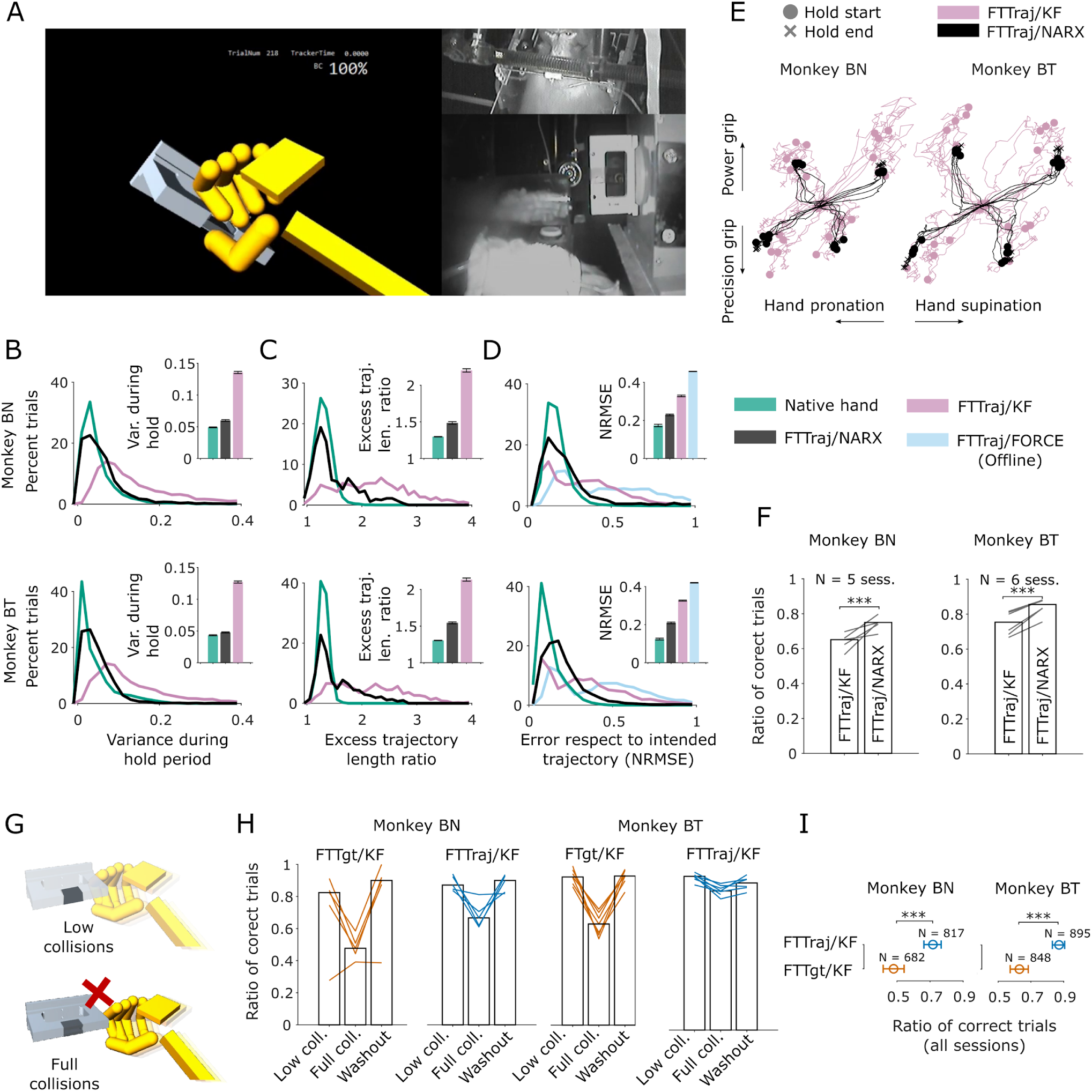
Performance of the training strategy. (A) Example video frame during a BCI precision grip using fit-to-trajectory combined with the NARX decoder (FTTraj/NARX, monkey BT). (B-D) Comparison of our full strategy to native hand grips and variants of the method. (B) Distribution of how much joint angles vary during the 500ms hold period in relation to the intended hold configuration in a trial. (C) Distribution of the ratio of executed to intended trajectory length for each trial. (D) Distribution of normalized root mean square error (NRMSE) of the attempted joint trajectory for each trial (compared to intended). Offline output of the FORCE RNN is included using the same trials of FTTraj/NARX (B-D) Number of trials/sessions: monkey BN (native 2454/6, FTTraj/NARX 756/5, FTTraj/KF 1492/5, FTTraj/FORCE 756/5) and monkey BT (trials/sessions: native 1142/6, FTTraj/NARX 1121/6, FTTraj/KF 1186/6, FTTraj/FORCE 1121/6,). Inset bars represent the median and standard error. (E) Example single trials using our full strategy and the KF-only variant (DOFs before DimExp). Trajectories start at the center (no grip) and end on the 4 hold points, corresponding to 40° vs. 180° handle orientation and power vs. precision grip conditions (circles and x’s). (F) Session performance comparison using our full strategy and the KF-Only variant. Bars: median performance. Lines: individual session performance. Trial performance distributions are significantly different (p<0.001, Clopper-Pearson estimation for the Bernoulli process). Same sessions and trials as in (B-D). (G) Illustration of the Low-collisions/Full-collisions task. Low-collisions: the virtual hand can go through the handle and achieve correct grips just by matching the right pose. Full-collisions: the hand would collide with the handle if not properly prepared for grasping. (H) Performance on the task before collisions were enabled (low-collisions), when collisions were enabled (full-collisions), and when collisions were disabled back (washout). FTTgt/KF: KF decoder fit with an “aim to target” strategy. FTTraj/KF: KF decoder fit with our trajectory following strategy. Bars: median performance across sessions. Lines: percentage of correct trials for each part of the task per session (N=5 sessions of monkey BN and 7 sessions of monkey BT). (I) Comparison of the ratio of correct trials during the full collision task (all trials, median and 0.999 CI, Clopper-Pearson estimation for the Bernoulli process).

Performance of our full strategy was superior to KF alone. We compared sessions using our full strategy (FTTraj/NARX) to sessions where subjects controlled the BCI only using the KF (FTTraj/KF, both decoders tested the same day). Hold variance, and trajectory length excess and error were lower using FTTraj/NARX when compared to the KF (Fig. 2B-D), and as visualized in example BCI trajectories (Fig. 2E). Overall task performance using FTTraj/NARX was also higher (Fig. 2F, Fig. S4A), with a well balanced distribution for both grip types (Fig. S4B). Output of the NARX layer was also better than just smoothing the KF output (Fig. S4C-E). Notably, the addition of the NARX layer did not require extra training by the subject, which gained performance from day-zero using FTTraj/NARX (Fig. S3A, Table S2).

Our strategy was also advantageous over a fit-to-target (FTTgt) approach^37^. Our virtual environment allowed two collision modes: a “low-collisions” mode were only the fingertips and the precision button were collidable, and a “full-collisions” mode, were the handle, fingers, and dorsum were collidable (Fig. 2G). While there were no performance differences after both subjects were proficient in FTTgt and FTTraj under low-collisions (Fig. S4G), FTTraj was advantageous when the collision mode was introduced to both subjects (Fig. 2H,I). This advantage remained even after subjects had more experience with the collision task (Fig. S4I, but see also Supplemental notes for caveats of this interpretation).

Our decoding approach was also superior to a state-of-the art autoregressive velocity-based method^43^. To tackle the problem of multi-dimensional grasping, we attempted to use a recurrent neural network (RNN) FORCE decoder^43^. Pilot studies demonstrated FTTraj/KF decoders were superior to FORCE, which exhibited poor control capacity and low trial performance (Fig. S3B, Monkey BT). The introduction of FTTraj decoders boosted performance from the start of training (Fig. S3B, Table S2). Output of the FORCE decoder produced highly variable and unstable trajectories that made holding the target hand pose difficult, a sign typical of poor velocity control. FORCE decoder output after finishing training Phase(iii) with FTTraj, when neural activity should have been better differentiated, was still deficient (Fig. 2D, Fig. S4F, offline test).

In sum, these results show our training protocol allowed monkeys to control grasping with high accuracy, and our approach had advantages over state-of-the art velocity-based decoders, and intention estimation techniques.

### Posture was decoded with greater accuracy than velocity, reflecting native grasping

Multiple studies have shown that posture can be decoded with more accuracy than velocity during native grasping^14,16,23–25^. To confirm that this was also the case during BCI use, we studied the contribution of position and velocity signals to online decoding in six sessions after training of Phase (ii) (performance > 80%, full BCI control). To measure this contribution, we re-ran the online decoders to predict independently the target zero-order and first-order kinematics and measured the correlation and fit quality by comparing them to the target trajectories. Neural activity predicted positions more accurately than velocities when looking at single trial prediction averages (Fig. 3A), and single trial prediction distributions for combined (Fig. S5B), and individual cortical areas (Fig. S5B), indicating that neural activity contained more position-related than velocity-related information during BCI control^16,23,14,25,36^ (see Fig. S5A for example predictions).

**Figure 3.**
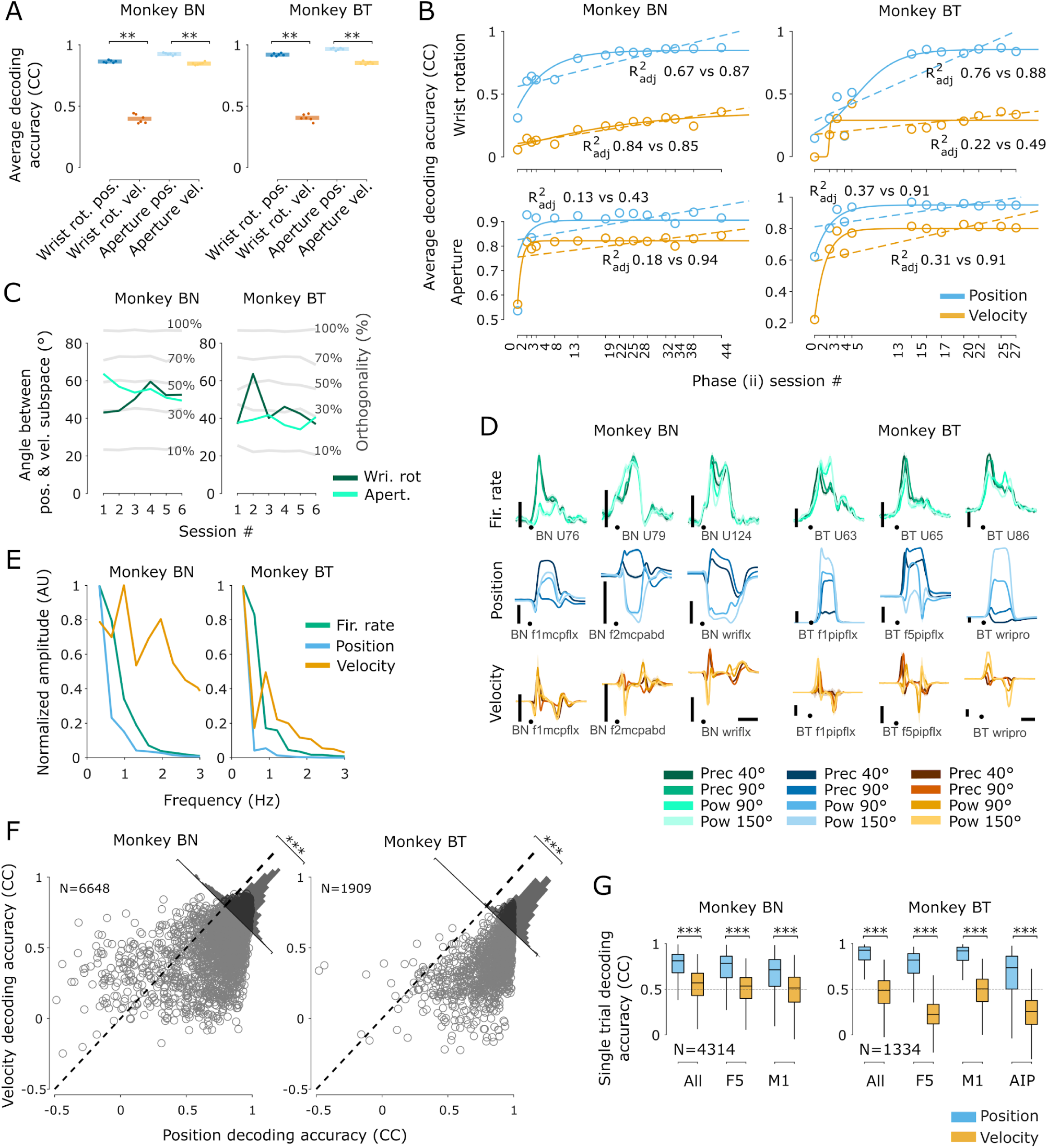
Neural activity correlation to position and velocity. (A,B) Decoding accuracy of kinematic variables during BCI control (CC: Pearson’s correlation coefficient). (A) Session average accuracies for 6 sessions after completing Phase (ii) of the training. We obtained CCs by comparing each decoded kinematic to the intended kinematic on each trial. Points: average CC of single-trial decodings for each kinematic. Bars: average of averages. **p<0.01, Wilcoxon signed-rank test on the averages. (B) Session average accuracies during learning of Phase (ii) (sessions previous to (A)). Lines: linear and sigmoid fits. Text: adjusted R² goodness-of-fit (linear vs. sigmoid). Accuracies were calculated as in (A). (C) Orthogonality between the neural population position and velocity subspaces for the wrist rotation and aperture variables in all sessions of (A). Gray lines: angle between the wrist rotation position subspace and the same subspace with a % of individual neurons randomly shuffled. (D) PETHs of example units and kinematics during the native grasping task. Positions: joint angles. Velocities: joint angle velocities. Dot: movement onset. Shades: s.e. Bars: Fir. rate 50 Hz, Pos. 10°, Vel. 100°/s, Time 1s. (U: unit, f1: thumb, f2: index, f5: little, mcp: metacarpophalangeal, pip: proximal-interphalangeal, wri: wrist, flx: flexion, abd: abduction, pro: pronation). (E) Average of the Fourier transforms from all PETHs of one native arm recording of each monkey. (F) Position vs. velocity single trial decoding accuracy on one native arm recording. Position (32) and velocity (32) kinematic wer predicted from neural activity on single trials using two independent KFs. CC was obtained by comparing predictions to average kinematics. Each point corresponds to one trial and one kinematic. (G) Position vs. velocity single trial decoding accuracy on one native arm recording considering all or individual areas. Same data as in (F). (A,F,G) **p < 0.01,***p < 0.001, Wilcoxon signed-rank test.

To verify that the above observations were not spurious correlations between the neural and kinematic signals, we looked back to the training of Phase (ii) which spanned 45 (Monkey BN) and 28 (Monkey BT) sessions preceding the six sessions above. If decoding quality was spurious (i.e. an artifact of coincident BCI output and cortical activity time dynamics), we would expect these results are independent of training history. On these training sessions, we looked for good recording days and selected 12-14 relatively even-spaced sessions from each subject (>60% correct trial ratios & >30 trials/condition). Decoding accuracy exhibited change during Phase training (Fig. 3B), indicating that these correlations were not just an effect of coincident time dynamics. The changes were also positive (i.e. decoding quality increased), indicating that the increases in prediction quality were useful for control as the subject gained performance over sessions.

The above results reproduced the asymmetry in position and velocity decoding quality observed in previous studies^16,23,14,25,36^. To check this was not just a sampling or decoding deficiency in our approach, we controlled our procedures using surrogate data. Velocity-like signals might be considered “harder to catch” as they correspond to sharp transitions in the neural signal that might not be captured by the decoder. Importantly, linear decoders are based on combinations of patterns and do not necessarily need these sharp transitions to build velocity outputs (Fig. S5C,D). To measure the capacity of the decoder to recover these signals, we built an encoding model where two tunable parameters (α, β) controlled the amount of position and velocity information encoded in artificial spikes (Fig. S5E, kinematics from 1 Phase (ii) recording per subject). Matching simulations to real spikes (Methods), we found that both, recordings with high-velocity content (high-β), and low-velocity (low-β) content were possible (Fig. S5F). Decoder tests revealed information was decodable in both regimes, but only one regime matched our case (Fig. S5G).

Previous results indicate that neurons in hand areas can contain both positions and velocity in orthogonal codes^15^. Tuning orthogonality is interesting as it is an indirect measure of decorrelation (i.e. independence) between the position and velocity codes, and if high, it indicates neurons can be multiplexed to help accomplish two decoding tasks^15^. To explore this in our BCI data, we compared the neural contributions to each degree of freedom (DOF) as determined by the decoder subspaces (the rows for position and velocity of the KF 𝐊 matrix, Methods). Our data indicated that for both monkeys, roughly 30-50% of the units had a different tuning for position and velocity control of the wrist (Fig. 3C) signaling orthogonality of the representations.

To further verify that the decoding results above are a property of the circuit and not an artifact of BCI use, we looked at recordings of the native hand task before BCI training started. We would expect the asymmetry found in BCI position and velocity would be similar for native arm kinematics. Example single unit peri-event time histograms (PETHs) exhibited a mixture of low and high-frequency components, resembling typical single joint angle position and velocity kinematics (Fig. 3D), but notably, neural signal peaks were not as sharp as velocity peaks. Normalized population averages of the frequency components of units and kinematics confirmed this observation, with units showing lower frequencies than velocity but not as low as position signals (Fig. 3E). This observation also extended to different brain areas (Fig. S6A,B, Supplemental notes).

We then decoded individual kinematics from single native hand trials using KFs. Accuracy results also reflected high position content (Fig. 3F), which was consistent for different areas (Fig. 3G), multiple sessions (Fig. S6C), and independent of whether the joint belonged to the hand or arm (Fig. S6D). The high ratio of position to velocity information was also confirmed through pairwise correlation of unit and kinematic average patterns. Even though this analysis misses the effects of linear combinations (in comparison to decoder regression, Fig. S5C,D), trial averages can better capture velocity-like sharp transitions which might be hidden by neural noise. On average, individual units tended to be more closely related to position than to velocity patterns (Fig. S6E and Fig. S6F). Accounting for potential pattern noise preserved these results (Fig. S6G, keeping only 90% of the variance through PCA^19,44^).

Overall, these findings indicate that the cortical posture signal was a major contributor to BCI control, and this control signal was orthogonal to velocity^15^. Notably, by using a position-velocity decoder, our strategy provided equal opportunities for neural correlates of position and velocity to evolve during Phase (ii) of the training. Yet, position correlates predominated, suggesting a preference by the grasping circuit to produce posture-related signals.

### BCI control performance was driven by enhanced posture information

The above results indicate there was a major role for posture information during control of our BCI. Given that cortical activity is the sole determinant of BCI output movements, neural representations of the required posture transitions must have been acquired to achieve proficient use. To confirm this, we looked for three neural signatures of this process during training of Phase (ii) (Fig. 3B): increase in overall population variance, increase in separation between conditions, and indications of change in population dimensionality during the movement phase (Fig. 4A).

**Figure 4.**
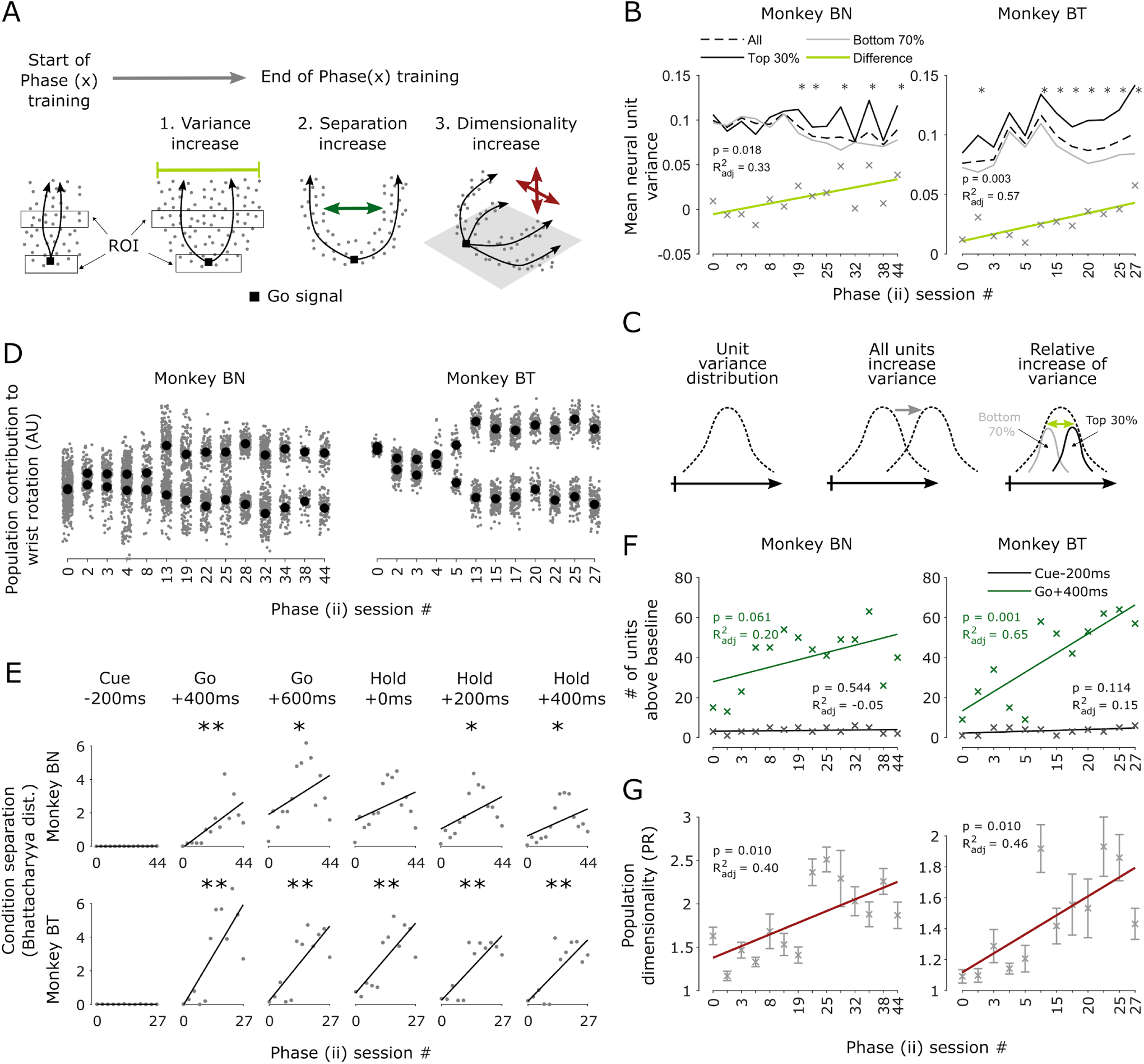
Changes in unit activity during training of Phase (ii) (A) Hypothesized changes in population activity from the start to the end of Phase training. Each point represents the combined activity of the neural population at a given point in time projected to a low-dimensional plane. Black arrows represent average neural trajectories for a given BCI movement condition (i.e. reaching the 40 or 180° angle). For analyses, we focused on activity in different regions of interest (ROI) during the trial. Although neural activity is represented here in low dimensions, we analyzed the full, high-dimensional activity. (B) Change in firing rate variance over sessions. Regression line was obtained from the differences between the mean firing rate variance of the top and the bottom contributing units. Units were ranked in the top-30% or bottom-70% according to their contribution to the Kalman filter output. ROI: 400 ms window around t=Go+400 ms. (*p <0.1, see Fig. S7A, total N c.a. 150 units/session) (C) Illustration of two potential changes in firing rate variance over sessions. Increase in the initial variance (left) can happen for the complete population (middle, all units increase variance) or for some of the units (right, relative increase of variance) (D-E) Separation of neural patterns over sessions according to their contribution to decoder output. (D) Each gray point represents the combined population activity contribution to the decoded wrist rotation output for one trial (position) in the units of the wrist rotation axis (Fig. S1). Black points represent the trial average for the power 40° (top) and power 180° (bottom) grasps. ROI: 20ms window around t=Go+600 ms for BN and t=Hold+300 ms for BT. (E) Separation between the clusters in (D) but now for different periods in the trial. *p<0.1, **p<0.01, p-value is for the regression line. (F) Increase in neural pattern separation over sessions. x’s represent the number of units whose difference in average firing rate (power 180° minus power 40°) is above the before-cue baseline. Green: 200 ms ROI around t=Go+400 ms. Black: 200ms ROI around t=Cue-200 ms (200 ms before cue start). Baseline: 97th percentile of the before cue period distribution of differences. (G) Increase in dimensionality according to the participation ratio (PR, ROI: entire period from the Go signal until the start of the grasp) (B,D-G) Changes were measured in the same sessions as in Fig. 3B. (B,F,G) Numbers indicate regression p-value and adjusted R². High p indicates that session # increase does not have a significant effect on the variable of interest.

To detect changes in variance, we measured the variance in firing rate of each unit used by the BCI and put together all the measurements to form a distribution. When taking all units together, session variance did not show a consistent trend over sessions (Fig. 4B, dashed line). However, ranking units by their contribution to the KF (weighted by entries of matrix 𝐊 in Eq. 1), we found a significant slope in variance for the top-30% contributing units relative to the variance in the remaining 70% (Fig. 4B, solid green line), and this difference became more significant towards the end of Phase training (Fig. S7A). Other splits (top < 30%) did not show a significant slope change relative to the remaining population. While the overall population variance remained constant, we observed an increase in intra-population variability (Fig. 4C), a finding consistent with previous work^45^.

To find pattern separation in the population, we looked first at changes in the neural drive 𝐊𝐳_𝑘_ in the KF (Eq. 1, Methods). This vector represents a low dimensional projection of the neural population contributing to each DOF before the assisting procedure is applied (Eq. 2, Methods), and thus, a proxy measure of pattern change^46^. The position kinematic contribution to the decoder output separated over training sessions of Phase (ii) for the wrist during the movement period (Fig. 4D), and this separation was consistent over different parts of the trial (Fig. 4E).

To measure these changes, we calculated the difference in firing rate of all units to form a distribution on each session (Methods). We used the pre-cue distribution as a baseline and compared it to the distribution after the “go” signal. Even though we could not find a significant trend in the increase of separation in the population as a whole (i.e. the median “go” separation did not increase over days relative to “cue”, Fig. S7B), we found a significant trend in the number of units whose difference was above the pre-cue baseline (Fig. 4F), indicating that changes were driven by an increasing number of units separating, and not gradual separation of the entire population. Per unit rate differences correlated to a good degree with the weights the 𝐊 matrix gave to each unit (Fig. S7C), a strong indication that these separations were indeed helping the decoder produce different trajectories.

We then looked into dimensionality changes in the neuronal population. The design of our task required subjects to progressively differentiate between conditions i.e., learn to separate the 40° and 180° grip, then, the power and precision grip. This should have manifested as increase in the capacity of the population to produce the different states when required. For position control, this implies redistribution of variance along dimensions in the neural population. We found traces of these changes when applying principal components analysis to the population after adjusting for asymmetries in population variance (Methods). A simple metric, the difference between the explained variance of PC1 and PC2, indicated that there was a redistribution in variance over sessions (Fig. S7D). A more formal dimensionality metric, the Participation Ratio^47^, also pointed to an increase in dimensionality over sessions (Fig. 4G).

Although we did not find a consistent trend in variance change during training (Fig. S7F), we found significant separation and dimensionality increase for both monkeys when learning Phase (iii) of the task (Fig. S7G,H).

Taken together, these results show that cortical activity encoded the required posture transitions during learning to use the BCI, and thus posture activity supports accurate hand BCI control for multiple degrees of freedom.

### Generalizability of our fitting approach

As our approach introduces a novel fitting procedure, we explored next its generalization properties^37^. Poor generalization on velocity-based decoders manifests as directional biases, i.e. the inability by the subject to adjust the movement in some directions (Fig. 5A). Although detrimental, the user can compensate for some of these biases by moving in non-biased directions, or moving at lower speeds. Poor generalization on position-based decoders manifests as the inability by the subject to reach specific regions of the workspace, a particularly detrimental feature (Fig. 5B,C). We explored two potential limits to generalization: limitations caused by the decoding strategy, and limitations caused by neural constraints.

**Figure 5.**
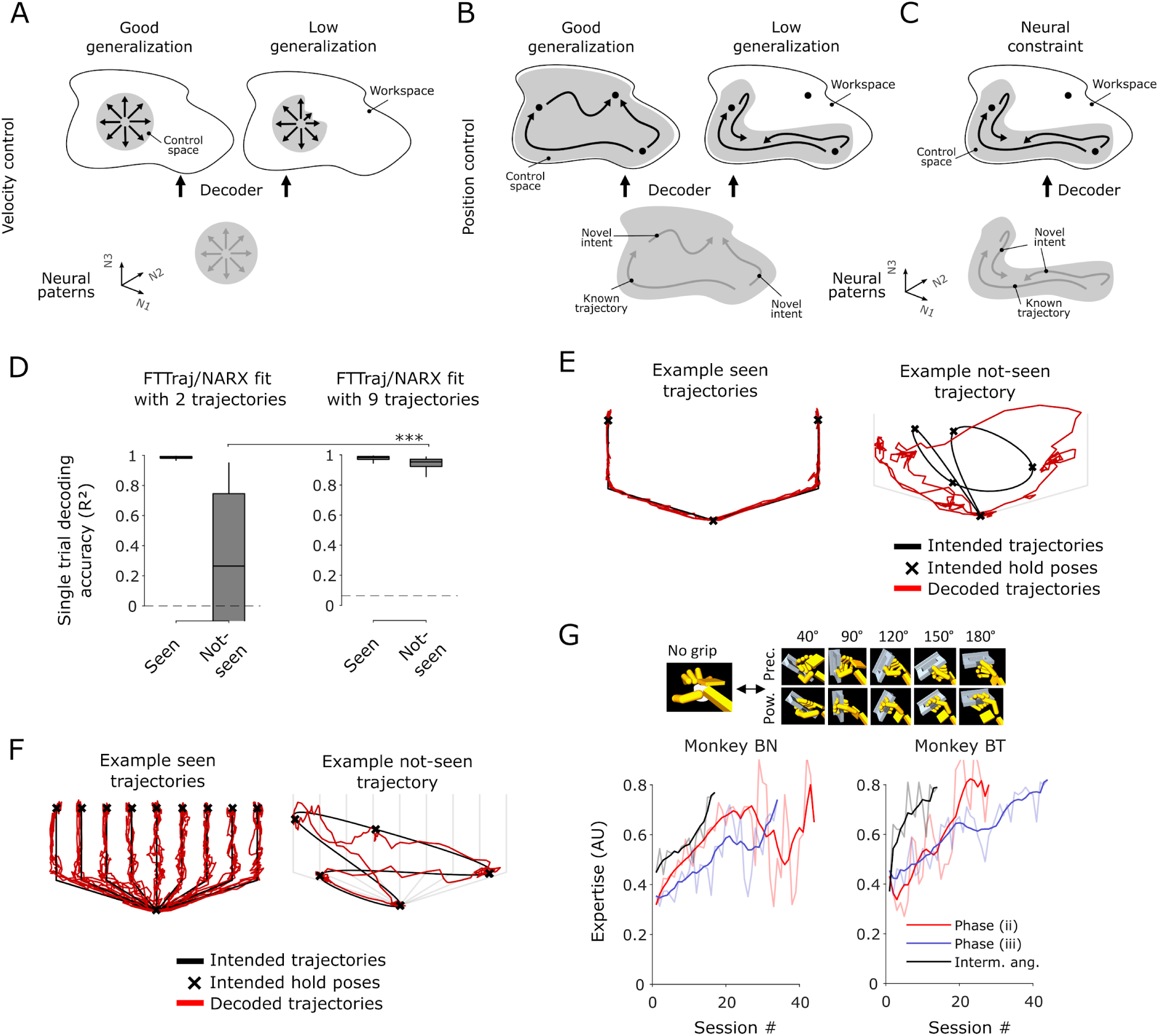
Properties of the fitting strategy. (A-C) Schematic of generalization properties for a BCI decoder. Same format as in Fig. 1A,B. (A) For a velocity decoder, poor generalization manifests as incapacity of the device to correctly exert velocity in particular directions. (B) For a position decoder, poor generalization manifests as incapacity of the device to reach parts of the workspace despite user intent. (C) Another cause for poor generalization are constraints in cortical activity failing to reflect user intent. (D) Effect of the number of trajectories used to fit the decoder. Left: FTTraj/NARX accuracy if 2 trajectories are used. Right: FTTraj/NARX accuracy if 9 trajectories are used. Seen: decoder accuracy for trajectories used to fit the decoder (N=450 trials). Not-seen: decoder accuracy for trajectories not previously seen by the decoder (N=500 trials, ***p < 0.001 Wilcoxon rank-sum test). Boxes: median, 25th, and 75th percentiles, and range of the distribution. Dashed lines: chance levels (Methods). (E) Example FTTraj/NARX decoder output for 2 fitting trajectories. (F) Example FTTraj/NARX decoder output for 9 fitting trajectories (E-F) Hold poses are points where the trajectories remain steady for 500ms. (G) Learning the intermediate angles task. Top: handle and grip configurations trained. Bottom: learning curves to achieve “Expertise” (ratio of correct trials times the average BC level on a session).

To explore limitations of the decoding strategy, we performed simulations using an RNN model of grasping^48^ combined with our full fitting procedure (FTTraj/NARX, Fig. S8A, Methods). To test generalization, we fit the decoder using a known set of movements (trajectories “Seen” by the decoder during fitting), and tested it by asking the decoder to produce a set of novel user intentions within the workspace (trajectories “Not-seen” by the decoder). Low generalization can be measured by the incapacity of the decoder to correctly reproduce novel intents from neural activity (Fig. 5B).

We started by fitting the decoders using 2 well spread trajectories in a 2D workspace. Although FTTraj/NARX decoders were able to produce the not-seen trajectories above chance levels when measuring accuracy (Fig. 5D), and below chance levels when measuring error (Fig. S8B), accuracy was low for not-seen trajectories. Decoders were able to follow the novel trajectories but movement was noisy and precision was low on hold poses (Fig. 5E), indicating overfitting. To check if more “seen” trajectories improved generalization, we increased the number of fitting samples to 9 equally distributed trajectories within the workspace (Fig. 5F). Accuracy and error of the non-seen trajectories significantly improved in this case (Fig. 5D, Fig. S8B), and decoder output was less variable, specially at holding poses (Fig. 5F). These results indicate that at least in 2D, our strategy generalizes, but well spaced within-workspace trajectories need to be considered during decoder fitting.

Given this restriction, would our strategy generalize to higher dimensions? As the number of dimensions increases, would the number of required intermediate trajectories exponentially explode? Simulating higher dimensions (Methods), we found that only a relatively small set of well spaced trajectories was required for comparable performance in seen and not-seen trajectories: 18 trajectories for 6 dimensions (Fig. S8C) and 27 trajectories for 12 dimensions (Fig. S8D).

Similar to limitations caused by the decoder, limitations caused by neural constraints manifest as inability to reach certain areas in the workspace. However, in this case restrictions are caused by restricted neural patterns^49,50^, which could be caused by over-training (Fig. 5C). To check if our Phase training and fitting approach could limit generalization by overtraining, we further trained our monkeys to produce novel postures corresponding to the intermediate angle configurations of the task (90°,120°,150° using power and precision grip) after finishing Phase (iii) of the training (Fig. S1, Fig. S3A). The trajectories used resembled the intermediate trajectories used in the simulations above. Both subjects were able to achieve our expertise criteria (BC level x performance > 0.8, Methods) in the intermediate angles after 17 (monkey BN) and 14 (monkey BT) sessions (Fig. 5G, Fig. S8E). As a comparison, it took monkeys ∼30-45 sessions to achieve the same performance criteria in the previous phases of training (Fig. 5G). This further demonstrates the extensibility of our approach to achieve novel posture configurations.

## Discussion

Despite reports of abundant posture encoding in hand-related areas during grasping^16,23,14,35,24,25,22,36,26^, the use of this cortical signal has not been verified for real-time control of a hand neuroprosthesis. Due to its inherent suitability for controlling effector trajectories and reported superiority in reach tasks^30–32^, velocity-based control has been a prevalent approach in BCI research. However, posture might be an equally informative signal for hand control^51,24,25^. Posture commands can be generally expressed as trajectories in the workspace of the effector (Fig. 1B). Our trajectory based fitting protocol enabled continuous prosthetic hand control with selection of more than one grip type at high accuracy. Posture could be better predicted from native grips and this finding generalized to our online decoder, which fitted both position and velocity. Offline analyses showed that the neural population exhibited signatures of variability increase, pattern separation, and dimensionality increase matching the zero-order structure of the task state-transitions, and hence encoded posture. Position control can suffer from overfitting to the specified trajectories, but we showed that our protocol can generalize, and with extended subject training, it does not limit the neural population to specific trajectory patterns. In summary, our findings show in a causal brain-computer interface setting^52,53^ that posture-related activity in the grasping circuit can effectively manipulate a prosthesis.

### Encoding of kinematic parameters in cortex

The differences in position and velocity decoding were consistent among the three implanted areas and the two task types. Recording from M1 during a rich, naturalistic task, Vargas-Irwin et al. found that a joint position decoder could describe the captured kinematics with high accuracy and explicitly decoding velocities did not improve performance^16^. Decoding from M1 and PMv in a related task, Bansal et al. found similar^36^, or better^23^ decoding qualities for position variables, and these differences were consistent across areas^23^. Okorokova et al. also found higher decoding accuracy for position in M1 and somatosensory cortex. The work from Aggarwal et al. found high correlations in decoding position from M1, PMd, and PMv, but found no noticeable differences in decoding velocity^14^. Our results align with the former studies, demonstrating superior decodability of position variables from hand-related areas. Interestingly, we also found good posture decodability for arm joints (Fig. S6D), countering the view that arm parameters tend to be represented as motion direction^29,27,54,55,28^. We note however, that similar results were found in the studies above while decoding shoulder, elbow, wrist, joint angles and 3D locations^16,23,25^, indicating that more sophisticated models to elucidate the encoding of movement parameters during naturalistic movement are required^56,57,24^.

Could there be other explanations for the position-velocity asymmetry in our results? The observed asymmetry in decoding performance may be attributable to our fitting protocol, which employed target trajectories to fit the decoders. However, it is important to note that we used a position-velocity decoder which at every session regressed neural activity to position and velocity, giving equal opportunity to both neural signals to increase correlation to any of the target kinematic patterns. In our decoder, the state transition 𝐀 matrix in the KF^58,59^ always had equal weights for the position and velocity states and the velocity values always had influence over the output (integrated velocity was combined with position). Fig. 3B illustrates the evolution of decoding accuracy for both position and velocity during Phase training. While position decoding accuracy improved steadily, velocity decoding progress was slower, suggesting a potential intrinsic limitation of the neural circuit in generating velocity patterns.

We found further evidence that a salient posture-based signal is represented in hand-related areas during grasping actions. Consensus regarding the representation of movement in the motor cortex, i.e. between low-level (muscle dynamics) or high-level (kinematics)^60^ features, or whether these signals cause or are just correlates of movement^61^, is not fully established. Previous reports of a salient position code are based in offline decoding^14,16,22–26^, and thus only offer correlative evidence of this property. As an online BCI study, our results suggest there might be a causal relationship^52,53^ between these codes, i.e. hand grasping areas preferentially encode posture. Prosthetic brain control requires correlates in neural activity that can produce a desired set of commands, and that has been usually kinematics^6–9,11,12^. Hence, the focus of our work is limited to the demonstrating these kinematic correlates are useful to control a prosthesis. Future studies where position- and velocity-based training protocols are tested in independent subjects are required to fully confirm this circuit property.

Our results suggest that the position and velocity decoding subspaces had differences in tuning, indicating that these representations had a level of orthogonality (Fig. 3C). Nason et al. demonstrated 2D individual finger control (index and combined middle, ring, and small) using velocity-only decoding^15^. Despite using a velocity-only decoder, their offline analyses indicated that linear decoders were capable of effectively differentiating between position and velocity correlates within their dataset^15^, suggesting that position and velocity had orthogonal codes, and hence position control was a viable avenue for future research. Work from the same lab also showed that the inclusion of position as a tuned parameter increased performance offline^51^. Our results confirm this prediction, and show the viability of position based control for a hand BCI.

Our training and fitting strategy combined position and velocity control. Nason et al. predicted that the addition of a semi-orthogonal position code could improve decoding quality^15^. However, combining position and velocity is untypical for recent approaches^37,43,7,15,8^. Do we have additional evidence that controlling with this dual code was better than controlling only velocity or only position? Our results using a FORCE decoder, indicate that combined position-velocity decoding was instantly advantageous over velocity-only (Fig. S3B). In addition, use of the second ReFIT innovation (which reduces the KF to velocity decoding), also caused performance drops (Fig. S3C). Our offline analyses also suggests that a velocity-only decoder would have been at a disadvantage with position-velocity decoding (Fig. 3). We did not explicitly test whether a position-only decoder was better than position-velocity decoder. However, the orthogonality results above were obtained directly from the KF parameters, indicating the decoder profited from this orthogonality in the code. At least offline, the addition of more information to the decoder, either by combining areas^59^, or codes^51^ usually increased performance.

### Practical implications for BCI development

We showed that position was a large driver of control in the BCI. What are the implications of controlling with position? The addition of position control to velocity control has been deemed detrimental^37^, but it was later shown this was only true when the subject moved simultaneously^62^, which is not the case in our experiment.

Position control requires activity patterns that cover the entire task workspace (Fig. 1B, Fig. 5B). Velocity control might be considered to be more flexible, as it only requires activity patterns to be available for each possible direction of movement (Fig. 1A), and if those patterns exist, and the subject can recall them, it is in principle possible to achieve any position in the workspace. However, the requirements for true, high-quality control are equivalent for both types of controller: to be able to achieve any point in the control space and to be able to remain at this point for any given amount of time. This is independent of whether the control space is spherical or irregular (Fig. 1A,B). This implies, e.g., being able to hold any direction of movement at constant speed during velocity control, or remaining in a particular point of the workspace for an indefinite time in position control. At this level, both types of control are equally optimal. The most likely limitation is the capacity of neurons to produce these patterns. For instance, it has been shown that neural activity restricts speed modulation during velocity control^37,63–65^. Our work shows that position control is possible for a state-based task such as grasping, further demonstrating the capacity of the circuit to produce arbitrary posture-based shapes^52^.

Our results confirm observations of a position code for grasping and demonstrate this code can be used online to control a prosthesis. Using this code for control assumes a static relationship between the neural signal and the decoder output (i.e., the output of the KF is largely determined by the neural code and fixed decoder parameters). Yet, previous work emphasizes that online dynamics can have a significant impact in output performance and might explain the deficiencies of position-velocity decoders respect to velocity^32,66^. These works suggest that performance deficits in position-velocity KFs could be attributed to biases caused by attractor dynamics^66^, and difficulties by the subject to conceptualize the physics of position-velocity control^32^. In contrast to these observations, our results suggest that position-velocity control was a better scheme than velocity alone. Although we did not systematically look for output biases, there was no evidence of such effects in the decoder output at any point during Phase training. Our observations also indicate that position control was more adequate (Fig. S3B,C), as velocity based control was ineffective at achieving targets, and the BCI was hard to maneuver and “floated around” without stopping. We argue that such dynamic effects depend on the natural position or velocity information available to control as hypothesized by Gowda et al. (p. 917)^66^. However, more systematic tests are required to verify this hypothesis.

Our subjects were able to achieve high accuracy in controlling 3 DOFs, including the selection of two grip types. Training of the second and third DOF took respectively 45 and 36 sessions for monkey BN and 28 and 45 sessions for monkey BT. Training periods of 30-70 days have been reported for low dimensional monkey tasks in past experiments^10,12,13^. A good comparison to humans performing a multi-dimensional artificial arm and hand task was reported for one subject, taking 14 days to learn 3D positioning of the hand, 8 extra days to master the closing of the hand, and an extra 20-30 days to master 3D hand rotation^7^. While our training times are consistent with previous monkey studies, they may be considered long for practical human applications. An important aspect to consider when comparing these times to human training is cognitive capacity. The limited cognitive capacity of monkeys compared to humans makes it difficult to convey the objectives of every training step. For example, our monkeys require 3-6 months to fully master the rotating handle task with the native arm, which will be a few hours of training for a human subject. Explicitly communicating task requirements to participants will likely accelerate training success within our framework. Yet, the training of high-accuracy tasks such as precision grasping might require longer times compared to tasks such as reaching.

Our method could be used in clinical applications by building datasets of manual tasks that explore a good subset of the hand configuration space. Our approach does not focus on the movement of individual joints^67,15^ but seeks instead to develop DOFs synergistically through tasks^7,17^. A property of our approach is that it is data-oriented and only requires trajectories of idealized kinematics in the workspace. Consequently, our method is potentially applicable to any movement that can be represented as a trajectory. These trajectories could be captured^20^ from non-disabled subjects doing the target task and replayed to subjects during Phase training. Our simulation results indicate that the trajectory-based approach works as long as a limited set of intermediate trajectories in the n-dimensional space are considered. Importantly, given the rich set of hand prehensile and non-prehensile actions, different training protocols beyond ours^7,17,67,15^ will likely be required to fully develop the natural set of hand skills.

### Links to BCI learning research

We observed neuronal activity changes over sessions of Phase (ii) and Phase (iii) of the task. Analyses of population activity change yielded two interesting results: 1) Although population variance was mostly preserved, the variance of neurons contributing to BCI output increased in relation to not contributing neurons (Fig. 4B,C); and 2) Pattern separation was not driven by gradual separation of the population activity, but by an increasing number of units separating (Fig. 4F, Fig. S7G). During BCI control, effector movements are causally linked to ensembles of neurons and these ensembles experience changes in their activity during learning^6,12,68^. These changes have been classified in fast, within session changes^49,46,69^, and slow, across session changes^45,50,69^. Studies suggest that during slow learning, the overall variability of units is preserved by increasing the variability of units contributing to the BCI while reducing variability on non-contributing units^45^. Additionally, slow changes are driven by gradual drifts in unit tuning^69^. Our results support this preservation of unit variability hypothesis, and indicate that the slow drifts happen unit-by-unit and not in the population at unison.

Acquiring proficiency in generating intermediate trajectories required several weeks of training for our subjects. Recent interest in the evolution of neural latent spaces in motor cortex^68,49,46,50,70^ has shown that neural patterns are constrained to a covariance space^49^ termed the “neural manifold”^49,44,50^. Under this framework, learning of novel pattern-to-effector mappings can occur within a session if the mapping exist within the covariance space^49,46^, or across days if the mapping is outside this space^50^. During Phase (iii) of the training, subjects learned the exterior trajectories of the workspace (Fig. S1D,E), and these trajectories were encoded in neural activity (Fig. 4). Under a linear transformation (essentially the KF), the intermediate trajectories are by definition inner parts of this 3D structure. Yet, neural activity did not cover this inner space within an intermediate-angle learning session. These findings suggest intriguing properties of the neural covariance space geometry that prompt further exploration.

We proposed an intention estimation approach based on trajectories instead of targets^37^. The performance increases observed in recalibration methods such as ReFIT^37^ come from the observation that neural activity during active control is different from that during passive observation^71^. Whereas previous results show these benefits can be seen intra-day^37,71^, w were more interested in the across-day effects of fitting to trajectories. Our results suggest that at this level, performance benefits are due to improved pattern separation, which might inform future BCI developers.

## Author contributions

AAT, JAM, WAS, and HS designed research. AAT, and JAM designed the algorithm. AAT, WAS, and HS performed research. AAT, and JAM analyzed data. AAT wrote the paper. AAT, JAM, WAS, HS reviewed the paper.

## Acknowledgments

We thank N. Bobb, S. Borchert, M. Dörge, A. Filippow, and K. Menz for technical and theoretical help. We thank F. R. Willett, and J. Goodman for helpful comments on an earlier version of this manuscript. This work was supported by Deutsche Forschungsgemeinschaft grants FOR-1847 B3, and SFB-889 C09, and EU Horizon 2020 project B-CRATOS GA 965044.

## Methods

### EXPERIMENTAL MODEL AND SUBJECT DETAILS

#### Animal model

Two adult rhesus monkeys (*Macaca mulatta*) participated in this study (monkey BN, male, 10 kg, 12 years old; and monkey BT, male, 16 kg, 13 years old). All animal housing, care, and experiments were conducted in accordance with the German and European laws governing animal welfare, and in agreement with the US Guidelines for the Care and Use of Mammals in Neuroscience and Behavioral Research (National Research Council, 2003), and the UK Guidelines for Non-human Primate Accommodation, Care and Use (NC3Rs, 2017). Authorization for conducting this experiment was delivered by the Animal Welfare Division of the Office for Consumer Protection and Food Safety of the State of Lower Saxony, Germany (permit no. 14/1442 and 19/3132).

### METHOD DETAILS

#### Experimental setup

We built an experimental setup in which monkeys could grasp a virtual rotating handle presented in a 3D graphics environment using a controllable virtual hand avatar. The virtual hand avatar was either controlled by the native hand or the BCI. During native hand control, the real hand of the monkey was tracked in real-time with a full arm tracking system, and information from the tracker was used to control the hand avatar and provide visual feedback. During BCI control, online neural or pre-recorded kinematic activity was used to drive the virtual hand. The native arm was occluded in all settings with a covering dark plate such that the animal could only receive visual feedback from the virtual display (Fig. S1A). The virtual hand and handle used a physics engine to detect virtual collisions (MuJoCo, DeepMind, UK). The virtual handle was carefully matched with an equivalent in size, and rotating, real handle such that the monkey received grasping feedback when using the native hand. Both the real and virtual handles rotated in synchrony and could be set in any orientation from 0° to 180°. We defined 0° to be the most leftward, and 90° to be the upright orientation of the handle (Fig. 1C). The real handle had sensors attached to detect either the power or precision grips. Equivalent physics engine sensors (pressure, crossing, and location detectors at different hand positions) were used to detect correct grips in the virtual environment. The task control programs were implemented using custom LabVIEW code (National Instruments, USA).

#### Task paradigm and native hand task

Animals were trained to sit in a primate chair and perform a self-paced delayed grasping task while looking at the virtual reality display. The monkeys could initiate a trial by placing their grasping hand (right hand for both animals) onto a rest sensor that enabled a red fixation dot on the middle of the virtual handle. After a random period (200 to 400 ms), the handle rotated from 90° to a random orientation (40°,90°,120°,150°,180°), and shortly after, a green color cue was shown next to the handle indicating either a power (left cue) or a precision (right cue) grip. After the grip type cue was turned off, monkeys had to withhold execution for a variable period (200 to 600 ms) until the red cue disappeared. The monkeys were then allowed to execute the grasp. A grasp was considered correct when the grip was held for 500 ms. Not completing the grasp within 2000 ms aborted the trial. All correctly executed trials were rewarded with a liquid reward after the hold period ended. Animals could then return to the hand rest position and initiate a new trial. Error trials were aborted and immediately followed by an error cue and error period. Only correct trials were used in the analyses. The non-grasping hand was gently restrained in a long acrylic tube. In the native hand task, we limited the handle orientations to 40°, 90°, and 150°.

#### BCI task

The BCI task was identical to the native hand task except for the control of the virtual arm, which was exerted by the computer, the animal (through neuronal activity), or a mixture of both. The native arm was unrestrained during the BCI task. However, lifting the native hand out of the hand rest sensor at any point during the trial invalidated it. During the BCI task, monkeys performed relatively small contraction movements while holding the hand at the hand rest sensor. These movements had only a very low correlation to the BCI task output, which we assessed in two recordings where monkey BT wore our high-accuracy hand tracker during the BCI task (Fig. S9). We did not measure EMG, so we cannot discard other effects muscle contractions could have had in the task (see also Supplemental notes for a discussion around this topic). We employed two levels of difficulty in the BCI task. In the “low collisions” mode, only the fingertips and sensors on the grasping handle that detected the grip type were enabled as colliding elements in the physics engine. In the “full collisions” mode, the full hand and grasping handle were enabled as colliding elements.

#### Native arm tracking

Previous to the recordings, monkeys were extensively trained in the native hand task while wearing an instrumented glove and observing their movements in the virtual environment. Finger, hand, and arm kinematics of the acting hand were tracked with eight magnetic sensor coils (Wave tracker, Northern Digital, Canada) placed onto the fingernails, the hand’s dorsum, and the wrist to compute the positions of the arm joints in real-time. This enabled tracking the positions of the 18 main arm and hand joints in 3D space including fingers, wrist, elbow, and shoulder with high accuracy (1 mm error)^72,59^. We developed a real-time inverse kinematics model to convert those in a 32 joint angles signal. The 32 joint angles were distributed on the shoulder (3), elbow (1), wrist (3), and fingers (25, 5 for each finger: flexion, spread, and rotation at the knuckle, and flexion of the proximal and distal interdigital joint). We obtained the joint angular velocity as the time derivative of each joint angle. The virtual arm model was built from exact measurements of each subject’s arm and hand dimensions and was driven by an equal number of joints (32 virtual servos).

#### Joint state-space and BCI workspace

In analogy to a “neural state-space” (i.e. a high dimensional space where each dimension is a neural unit), we refer to the space of possible combinations of the 28 hand joint angles (including wrist and fingers) as the hand’s “join state-space” or simply “joint space”. In the BCI we refer to the set of synergies we used to control as the “degrees of freedom” (DOF) which form the BCI “workspace” (as the term “workspace” in robotics).

We controlled three DOFs: grip aperture, wrist rotation, and grip selection. *Grip aperture* was a positive-only axis where position 0.0 indicated the hand is open (flat) and position 60.0 indicated that a grip is completed (either a power or a precision grip). The *wrist rotation* axis was aligned to the handle orientations creating an equal distribution for each grip target from the 40° precision to the 180° power. To compensate for the difference in wrist orientation between the power and precision grip, there was an offset of ca. 50° between the grips (e.g. wrist angle for precision 90° corresponding to power 40°). In the *grip selection* axis, 0.0 shaped the hand to a precision grip and 60.0 to a power grip. Position (0, 90° power, 30) in these three axes was the neutral hand rest position. To encourage the subjects to prepare the hand configuration in time, and following the native kinematics (Fig. S2), we designed the DOF space in such a way that curved trajectories were required to successfully prepare the hand before encountering the handle, which explains the bent trajectories in Fig. S1. Not following the trajectories would result in incorrect hand preparations and potential collisions with the handle (Fig. S4H). To match the native task visually (reach-to-grasp movement) we automated the hand reach movement and synchronized its progression with the grip aperture axis according to hand/arm psychophysics^73^. Note that we prefer the term “grip aperture” instead of “hand opening” as this DOF is slightly different for each grip type. Fore power grip, this DOF controls the closing of the index, middle, ring and small finger. For precision grip, this DOF additionally controls closing of the thumb.

Movement within the three variable DOF space was mapped to movement in the 28 joint state-space through a heuristic dimensionality expansion (DimExp) layer built from the full native hand dataset. The native hand dataset consisted of 10 different trajectories (power and precision grip, each with 5 different wrist orientations) in a 28-dimensional space. Each trajectory represented a state change from a fully open hand to a target grip type at one of the 5 target angles. Trajectory times were matched so that all trajectories had the same number of time points (T). Using the 5 power trajectories, we built a power grip state-space by smooth interpolation between time-matched points in the trajectories. This resulted in a T×K×28 tensor with T the number of time points, and K the number of interpolation steps between the left-most to the right-most angle (ca. 300). We used extrapolation at the extremes of the state-space hence preventing the 40° and 180° angles were easier to the subject by saturating to the extremes. To convert from DOF variables to the full power grip joint state-space, we linearly mapped the T index to the grip aperture axis and the K index to the wrist rotation angle. In this form, any point in the space (T, K) had a matching 28-dimensional point. We followed the same procedure to construct the precision grip state-space. To join the two spaces, we aligned the two state-spaces by angle (90° precision corresponding to roughly 40° power) and interpolated between the two spaces with L interpolation steps between grip types (ca. 60). This resulted in a T×K×L×28 tensor and (T, K, L) were linearly mapped to our three DOF variables.

#### Neural recording

After successful training in the native hand task, the animals were implanted with a titanium head holder attached to their skulls, and, in a subsequent surgery, with permanent floating microelectrode arrays (FMA, MicroProbes, USA). We targeted three key cortical areas of the grasping circuit: anterior intraparietal area (AIP), ventral premotor area F5, and the hand area of primary motor cortex (M1). In monkey BT we recorded from 6 individual 32-channel arrays implanted into AIP, F5, and M1. Monkey BN was implanted with 8 individual 32-channel arrays but only six arrays were used (F5 and M1, Fig. 1E). On each array, the lengths of the electrodes increased towards the sulcus and ranged from 1.5 (1st row) to 7.1 mm (4th row). Further surgical procedure details have been described in previous experiments^72^.

The neural signal was sampled at full bandwidth with a frequency of 24 kHz at 16 bits resolution, and simultaneously sorted online and saved to disk (RZ2 BioAmp Processor, Tucker Davis Technologies, USA). For online spike sorting, the signal was hardware filtered (bandpass 0.3-7 kHz) and spikes were detected using the RZ2 default hardware and software. Sorting was performed daily and manually on each channel, taking the previous recording session as a template. Inactive channels were identified by comparing their PETHs distributions at the beginning of the session to a uniform distribution (Kolmogorov–Smirnov test) and then discarded. No further selection criteria were applied to select the channels. All active units and multi-units were used for online decoding and offline analysis.

The number of recording sessions analyzed varied for each experimental task. For the analysis of the native hand task, we used data from 6 sessions of each animal. Each native hand recording session consisted of 481 ± 103 trials (mean ± SD) and 194 ± 2 single/multi-units for monkey BN, and 221 ± 85 trials and 189 ± 3 single/multi-units for monkey BT. The number of BCI sessions and unit counts varied for each BCI experiment with sessions ranging from 100 to 250 trials. The number of units and multi-units varied with counts ranging from 150 to 180 for monkey BN, and 130 to 160 for monkey BT. Primary motor, premotor, and parietal areas are all potential targets for motor prostheses^74,10,75,14,59,35^. All units and multi-units were analyzed and online-decoded together, irrespective of brain area. We refer to them simply as units.

#### Overview of the training approach

We incrementally trained^7,17,39^ three degrees of freedom (DOF) in three phases: (i) grip aperture control, (ii) wrist rotation control (hand pronation/supination), and (iii) grip type selection (Fig. 1F, Fig. S1, Fig. S2). The three phases were trained sequentially, each over several training days, not moving into the next phase until a performance criterion was met (>80% performance, Methods). To convert the 3 DOFs into high-dimensional hand joint values we used a dimensionality expansion (DimExp) layer in the decoder (Methods).

To fit the decoder, we used a three-stage procedure (Fig. 1G-J, Methods)^37,40,41^. In the observation stage (stage 0, Fig. 1G), the subject performed the delayed grasping task while observing automated grip attempts (intended transitions in DOF space). Neural activity and the intended transitions during observation were used to fit an initial KF decoder.

Next, subjects attempted the grips using neural activity (stage 1, Fig. 1H). Due to misalignments between the intended and effective neural activity, control performance in this stage was typically limited (attempted transitions in DOF space), and decoder re-fitting increased accuracy. Note that we did not explicitly measure these performance differences as there is well-known literature that shows improvement with intra-day recalibration^37,76,71^. Contrasting previous approaches, we then fitted a new KF decoder using a *fit-to-trajectory* (FTTraj) approach. This approach centers around transitions instead of targets^37^, attempting to preserve the desired position-based features (Fig. S1, Fig. S2). FTTraj was divided in two steps (Fig. 1I). In step 1, each grip transition attempt from stage 1 was projected to its corresponding intended transition (Methods). In step 2, the projected vectors were used to build a new adjusted trajectory for each attempt. Neural data from stage 1 and the projected trajectories were used to fit a new decoder (stage 2, Fig. 1J).

The KF is a linear state estimator that has proven useful in multiple BCI applications^37,9,77,71,15^. However, it is known the KF has limited accuracy when honing in on targets^43,37,63–65^. Non-linear maps can find idiosyncratic correlations between neural activity and kinematics or kinetics^78–80,65^, and thus, could improve target accuracy^43,65,81^. However, non-linear regression methods have an increased capacity to overfit, and in decoder context, make it harder to discern the properties of the driving neural activity. We found the best performance by combining the KF decoder with a non-linear recurrent network (non-linear autoregressive with exogenous inputs, NARX)^42^ which we applied at the end of each training phase (Fig. 1K,L).

#### BCI decoding

Spike counts and behavioral data were streamed from the RZ2 and a behavior control PC to a decoding PC using the UDP protocol. Custom LabVIEW code was used to store the online signal and run the decoders. MATLAB (Mathworks, USA) code was used to fit the decoders. The decoder was fed online with spike counts binned at 20ms without further processing. The main online and offline decoder was a position-velocity Kalman filter previously described^59^. The filter follows exactly the implementation of Wu et al.^58^. The filter had 3 to 7 state variables (1-3 positions, 1-3 velocities, and 1 offset term) depending on the number of DOF of the current training phase. Only the position state output of the decoder was used to drive the BCI. For the collision experiment, we implemented both innovations of the ReFIT protocol^37^, but found a substantial decrease in performance with the second innovation (Fig. S3C), and therefore used only the first innovation, i.e., we did not remove the position estimate from the filter.

#### Relating BCI neural activity to the Kalman filter output

We reinforce target trajectories over multiple sessions using a Kalman filter (KF)^58,59^. At its steady state, the KF takes the form:

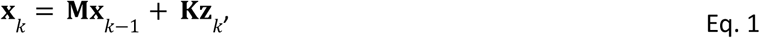

with 𝐱_𝑘_ the estimated kinematic state (position and velocity) at time bin 𝑘, 𝐳_𝑘_ the binned spike counts, and 𝐌 and 𝐊 two constant matrices. Neural input 𝐳_𝑘_ drives the filter and linearly affects the decoder output (𝐌𝐱_𝑘−1_ a non-linearity that acts as a smoothing filter on 𝐱_𝑘_). Having a linear drive 𝐊𝐳_𝑘_ implies that the neural population activity shape is related to the kinematic output shape 𝐱_𝑘_, which is a lower-dimensional projection of 𝐳_𝑘_ through 𝐊, i.e., neural activity should reflect velocity- and/or position-like characteristics (Fig. 1A,B).

#### NARX layer and FORCE decoder

To complement the Kalman filter, a non-linear autoregressive with exogenous inputs (NARX) recurrent network model was used (function “narxnet”, MATLAB Deep Learning Toolbox). The main difference between NARX and RNNs is that it receives feedback from its output and not its hidden states^42,82^. Our NARX had two layers: layer 1 was a tanh layer with 20 units, and layer 2 had 3 units (corresponding to the number of target DOFs). The network received as input the position output of the Kalman filter (3 states) and its output (3 states). For fitting, we used Bayesian regularization, and the Real-time Recurrent Learning algorithm (RTRL), an alternative to back-propagation through time that has gained recent interest due to its advantages in online settings^83^. Importantly, the network was fitted offline only once, using an example Kalman filter session as input, and the target trajectories as output. At the start of training of monkey BT, we also used a FORCE RNN decoder, which we implemented exactly as in^43^ using code shared to us by the authors (90% velocity control).

We used the hybrid scheme to improve the accuracy of the kinematic output. The output of the Kalman filter is fed to a two-layer recurrent neural network^42^ fitted to produce the target trajectories. This configuration was motivated by the principle of an “information bottleneck” (e.g., as in variational autoencoders^84,85^) in the context of hand state dimensionality reduction^86^. Following this principle, we first trained the subject to produce the output without the non-linear layer to ensure robust features were first developed. In this respect, the noisy output of the Kalman filter is advantageous, as it should pressure the brain to produce the correct patterns over training to achieve the basic target patterns. Once the basic patterns were developed, non-linear finetuning was added on top to improve performance.

Other practical factors motivated this arrangement: non-linearities on the transitions and hold states, potential of extensibility to more complex workspaces, and in our data, low tolerance to noise of pure non-linear/RNN solutions such as FORCE and NARX (see however^81^). A double decoder setup comes at the cost of complexity. We believe however this is justified for our application. The Kalman filter has been applied in numerous BCIs^37,9,77,71,15^ likely due to its simplicity, tolerance to neural noise, and compute cost, which is essentially two matrix multiplications once the filter reaches its steady state. Yet, characteristics of the neural data limit its accuracy^37,63,64^, requiring alternative or hybrid solutions^43,63–65^.

Notably, addition of the NARX layer required no extra training by the subject. In our experiments, we added this layer right after finishing training of Phase (ii), and the performance gain was immediate.

#### Incremental training of DOFs

To train degrees of freedom (DOFs) incrementally, we trained the subjects in three phases: (i) grip aperture control, (ii) wrist rotation control (hand pronation/supination), and (iii) grip type selection (Fig. 1F, Fig. S1). The three phases were trained sequentially, each over multiple training days, not moving into the next phase until a performance criterion was met. To train each phase, we followed a BCI assisting procedure in which a certain percentage of task control was done by the computer and the rest by the subject. The level of subject control increased until achieving full brain control (BC, level 0% to 100%). The criteria to move to the next phase was achieving BC=100% and task performance ≥ 80% over several days.

The choice of this sequence was motivated by saliency, design of the task, and native task training history. Grip aperture is the most basic DOF in grasping and for this reason, we chose it to be first. It was also lumped with the reach of the hand, making it a priority to achieve the hold configuration. Wrist rotation was the next, most salient DOF, as in our task the handle orientation is a large feature in the visual representation of the hand. The grip type is visually the least salient feature and for this reason, we chose to train it after the monkeys had more experience with the BCI. A similar sequence (wrist control before individual hand DOFs) was followed in related works^7,17^.

#### BCI training

Monkeys were trained to use the BCI through an assisted procedure in each task phase using the predefined intended transitions (Fig. 1). At time step 𝑘, this assistance had the form:

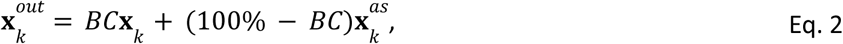

with 𝐱_𝑘_ the kinematics predicted by the Kalman filter from neural activity, 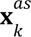 the pre-recorded kinematic transitions, 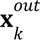 the output kinematics driving the virtual hand (before DimExp), and BC the brain control factor. The BC level variable was adjusted by the trainer from 0% to 100%, with 0% being fully assisted control (pre-recorded trajectories), and 100% full brain control. Intermediate values would mix proportionally pre-recorded trajectories and subject control. To calibrate the BCI, we followed a three-stage procedure to first initialize and then refine the decoder. At the start of each session, the monkey first observed the virtual arm perform 10 automated trials of each grasping condition (Observation or Stage 0). The model arm followed pre-defined latent trajectories and grasped the handle error-free. We used the neural data during this stage and the corresponding intended trajectories to fit an initial decoder. This decoder was used on a new stage (Initial or Stage 1) at variable BC levels (depending on the progress of the animal). The subject used the BCI until completing 10 correct trials of each grasping condition and a new decoder was fit using fit-to-trajectory (or fit-to-target on the collision experiments). The new decoder was used in Stage 2 (Fig. 1J). The NARX decoder could be alternatively enabled on Stage 2 (Fig. 1L). In our experience, more than 10 grips per condition or more than 3 stages did not significantly improve decoder performance. Importantly, all BCI data reported corresponds to BC 100% except for results showing learning curves and phase training.

#### Fit-to-trajectory procedure

To calibrate the decoder using fit-to-trajectory, the output position and velocity of the previous stage were replaced with the attempted positions and velocities projected on the intended trajectory. At hold points, positions are replaced by the intended target position, and velocity is set to zero. We calculated the projection (see Fig. 1, and Fig. S10A-D for a rationale) as follows: (a) Divide each attempted and intended trajectory into decoder time steps of equal time (20ms in our case) resulting in 𝑁*^att^* and 𝑁*^int^* segments. (b) Determine the segment length of the attempted onto intended trajectory as:

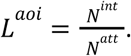

(c) Let the attempted and intended trajectory be defined as n-dimensional parametric curves 𝑓*^att^* (𝑡) for parameter 𝑡 = {0, 1,…, 𝑁^𝑎𝑡𝑡^} and 𝑓^𝑖𝑛𝑡^ (𝑡) for parameter 𝑡 = {0, 1,…, 𝑁^𝑖𝑛𝑡^}. Here, each value 𝑡 matches a trajectory time step. (d) The projection of attempted onto the intended trajectory is determined by finding the values 𝑓^𝑖𝑛𝑡^(𝑡) corresponding to parameters 𝑡 = {0, 𝐿^𝑎𝑜𝑖^, 2 × 𝐿^𝑎𝑜𝑖^,…, 𝑁^𝑎𝑡𝑡^× 𝐿^𝑎𝑜𝑖^}.

The values obtained at points 𝑡 = {0, 𝐿^𝑎𝑜𝑖^, 2 × 𝐿^𝑎𝑜𝑖^, …, 𝑁*^att^* × *𝐿^aoi^*} can be used to produce the position 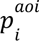 and velocity vectors 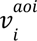 of the attempted onto the intended trajectory. After this, the Stage 1 neural activity, 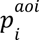, and 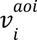 are used to fit the Stage 2 decoder. Note that this transform preserves the time taken in the attempted trajectory and hence matches the number of neural activity timesteps. We provide MATLAB code with an example implementation of this method in the supplemental section (see Table S1 and Fig. S10E,F).

### QUANTIFICATION AND STATISTICAL ANALYSIS

#### Decoder comparisons

To compare decoders online, we paired decoding blocks of a similar number of trials from each decoder type from the same recording day. The sequence of decoding blocks was chosen randomly each day. On the fit-to-trajectory and fit-to-target comparisons, each decoder was fit independently following our 3 stages approach. Unless noted, a decoding block usually consisted of ca. 100 trials. On the fit type comparison, the same decoder was used in the three blocks: partial collisions (100 trials), full collisions (100 trials), and washout (20 trials). For the comparisons to native hand accuracy, trial-averaged trajectories, and trial-averaged hold poses were used as the intended trajectories and target. Experience before online decoder comparison experiments is summarized in Table S2.

#### Comparison of neural and kinematic signals in the BCI and native tasks

To measure the amount of position and velocity information on the neural activity of BCI sessions we rerun every session offline using the same decoder used online and obtained the position and velocity outputs out of the KF state. We then calculated Pearson’s correlation coefficients (CC) or coefficient of determination (R²), in their standard form, on single trials by comparing the outputs to the intended trajectories on any given trial. Similar results were obtained by fitting independent position and velocity KF filters (using the first 20% of the session to fit). For single-trial unit and native joint angle comparisons, we used regression through KF. For offline regression analyses, we preferred the KF as a predictor as it was a closer comparison to the online setting, following previous works^16,59^, and because it performed better than other alternatives such as least-squares in our datasets. We then used CC or coefficient of determination (R²) comparing single decoded trials to matched average kinematics. All offline regression analyses used cross-validation using the initial portion of the recording (first c.a. 10 trials of each condition or first 10-20% of the session) to fit a decoder and predict the rest of the session. A larger number of training trials did not change results substantially. To compute PETHs in single unit and native joint angle comparisons both spike rates and kinematics were split in 20ms bins, averaged within conditions, and smoothed with a Gaussian kernel (50ms SD). Before averaging, we employed time-warping^87^ by aligning to trial events. For the Fourier transform analysis, we applied Fourier transforms to the PETHs, then normalized to the largest Fourier amplitude of each signal, and then averaged all results over units or kinematics.

#### RNN models

Owing to the intricacies of training monkeys to perform arbitrarily many hand poses within the BCI workspace (e.g. task the monkey to hold the hand in mid air), we used a recurrent neural network (RNN) model of grasp areas^48^ to test generalization of our approach. Using the RNN, we constructed neural state-spaces encoding 2D target kinematics and used the produced artificial neural activity to fit our decoder. Neural state-spaces were built by training randomly initialized RNNs to produce a set of target trajectories in the kinematic space (“seen trajectories”), and a set of random trajectories within the kinematic space (“not-seen trajectories”). We then used the RNN unit activity to build fit-to-trajectory NARX decoders (FTTraj/NARX), and then used the decoders to predict seen and not-seen trajectories (1 decoder per RNN).

In the simulations, we used single module RNNs with 20 units, a recttanh non-linearity, and α=1e-5, β=1e-2 as regularization parameters. RNNs were fed with latent neural signals and trained to produce the position **x** and velocity **v** kinematic (Fig. S8A). Signal **s** was produced by multiplying the normalized target latent position by a random NxM matrix with M the number of latent kinematics and N=1.5*M. Internal unit activations (**u**) were used to fit the decoders. We simulated 6 different settings: 4 in 2D, 1 in 6D, and 1 in 12D using artificially generated trials (90 trials for 2D, 180 trials for 6D, and 270 trials for 12D) with time dynamics similar to the real data (including 500ms hold periods). In the 2D simulations, we used artificial “seen” trajectories similar to the trajectories on our fitting paradigm. In the 6D and 12D simulations, “seen” trajectories were generated as curves within a hypersphere using natural cubic splines. For each setting, we generated 50 additional not-seen conditions using splines and 500ms hold periods. We trained 10 randomly initialized RNNs for each setting and used all trials to compare performance (50 total). For each RNN we trained a new decoder using 50% of the data and predicted the other 50%. Decoders were fitted with the internal activity of the RNN (**u**) and the target latent positions and velocities using fit-to-trajectory and NARX as in the online case. To train the RNNs, kinematics were injected with Gaussian noise of 30% of the signal amplitude, and unit activations with Gaussian noise of 15% of the signal amplitude. Chance levels were estimated by fitting decoders with neural data shifted by a random number of time bins.

#### Encoding model

To test the capacity of the Kalman filter to recover position and velocity information from binned signals, we encoded the kinematic variables of the BCI task as spike rates. To do this, the intended position and velocity signals of an example session from each animal (2 positions, 2 velocities, 20ms samples) were multiplied by scaling factors (α, β), Z-scored, injected with Gaussian noise (20% of the signal amplitude), and offset as positive only values. The resulting signal was then multiplied by an Nx4 normally distributed random matrix (N=130) and given to a Poisson filter to produce spike-like counts. A portion of the resulting artificial spike train (10%) was used to fit a Kalman predictor, and the resulting decoder was used to predict the kinematic variables of the remaining session. Decoding accuracy of the model data was measured as in the previous cases by comparing the output of the decoder to the intended kinematics. We ran each model parameter combination 10 times with different noise and random matrices and reported the average and the 95% confidence interval of each output (CCs or total spikes). We then used the total amount and spike count distribution of spikes generated by the model to match the number of spikes captured in the online session (a proxy measure of the total information available).

#### Orthogonality analysis

To measure orthogonality between the position and velocity subspace, we used the 𝐊 matrix used by the KF on each session online. Each row of 𝐊 represents one decoded dimension and each entry the weight the decoder gives to each neuron for a specific dimension. The rows of 𝐊 can be visualized as a vector in N-dimensional space (N the number of units) and the axis along that vector corresponds to the decoded variable subspace. To obtain the angle between the axes we used the MATLAB function “subspace”. To produce an estimate of the % of orthogonality we used a shuffling analysis. This % levels represent the angle between the wrist rotation position subspace (one 𝐊 row) and the same subspace with a given % of individual neurons randomly shuffled for a given session (same 𝐊 row but with a % of the row entries shuffled). We repeated the shuffling 100 times and reported the averages. In this metric 0% indicates no orthogonality, > 0% indicates orthogonality on a given % of dimensions, and 100% indicates full orthogonality. Similar results were obtained with normally distributed random vectors instead of shuffled neurons and for the aperture subspace (not-shown).

#### Analysis of changes in neural population activity

To perform analyses in population changes over sessions, we used the same neural activity and decoders used online during training of Phase (ii) and Phase (iii). For Phase (ii) we used the same recordings as in Fig. 3B. For Phase (iii) we used a well-spaced selection of recordings with good performance (around 80%). Unit activity was not further processed beyond time warping to obtain trial averages when needed (same procedures as in other analyses). Importantly, kinematic activity was obtained directly from the decoder output i.e. before applying the BC training help level.

##### Variance increase analysis

Variance of an individual unit was obtained from the combined firing rates during the 400 ms ROI of individual trials (20 bins per trial). The mean neural unit variance for a session was obtained as the mean of the individual unit variances. Trials in this analysis were not separated by condition. To categorize unit activity in the top 30% and bottom 70% we used the absolute value of the rows of the K matrix of the Kalman filter. We used the same KF obtained after fit-to-trajectoy calibration of each session. For Phase (ii) we used the row corresponding to the wrist rotation DOF and for Phase (iii) we used the row corresponding to the grip type DOF.

##### Pattern separation analysis

For the two 200 ms ROIs, we calculated the difference in firing rate using two methods: the mean difference and the Bhattacharyya distance. Differences in a session were computed independently for each unit. For a given unit, we used all the firing rate bins during the ROI of individual trials. Rates were separated by condition, forming two distributions for each unit (power 40° vs. power 180° in Phase (ii), and power vs. precision for Phase (iii)). The mean difference was obtained by taking the mean of each distribution, and then subtracting the mean of one condition from the other. This difference could be positive or negative but we use the absolute value. The Bhattacharyya distance was obtained by comparing each distribution directly (only positive values). Differences/distances for all units formed a distribution, one for the before-cue period (Cue - 200ms) and one for the after-go period (Go + 400 ms). The distribution for the before-cue activity was used to determine a “low-difference” baseline. This was a period where there was no firing rate separation between conditions but similar statistics (verified from PETHs of sample sessions). Differences below the 97th percentile were considered low. The Bhattacharyya distance is a measure of similarity between two probability distributions and extends the Mahalanobis distance to an entire distribution instead of just a sample.

##### Dimensionality analysis

We normalized population unit activity before obtaining the per session PCA and the participation ratio^47^. Firing rate normalizations are typical before applying PCA to deal with the sensitivity of PCA to high-variance (see for instance refs.^47,75^). Our normalization procedure was motivated by possible asymmetries in population variance that alter the dimensionality seen by the decoder (see also Fig. S7E for a rationale). To obtain the normalized neural activity matrix 𝐙*_N_*, we applied the following formulas:

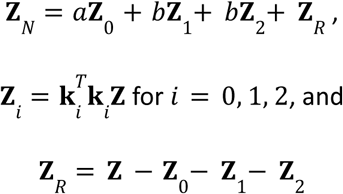

Here 𝐙 is the binned spike activity matrix over time (each row a unit); 𝐤_𝑖_ the normalized i-th row of the Kalman matrix 𝐊; and 𝑎, 𝑏, 𝑐 the scaling factors for the wrist rotation, hand close, and grip type DOFs. The procedure can be visualized as an N-dimensional transformation that scales the “cloud of points” in 𝐙 along the 1D axis determined by each 𝐤_𝑖_. Factors 𝑎, 𝑏, 𝑐 were obtained as ratios from the BCI workspace such that the workspace formed an equilateral cube. Factors 𝑎, 𝑏, 𝑐 were the same for all sessions. The participation ratio was calculated exactly as in ref.^47^.

