## Supplemental material for "Accurate neural control of a hand prosthesis by posture-related activity in the primate grasping circuit"

### Supplemental information

#### Supplemental figures

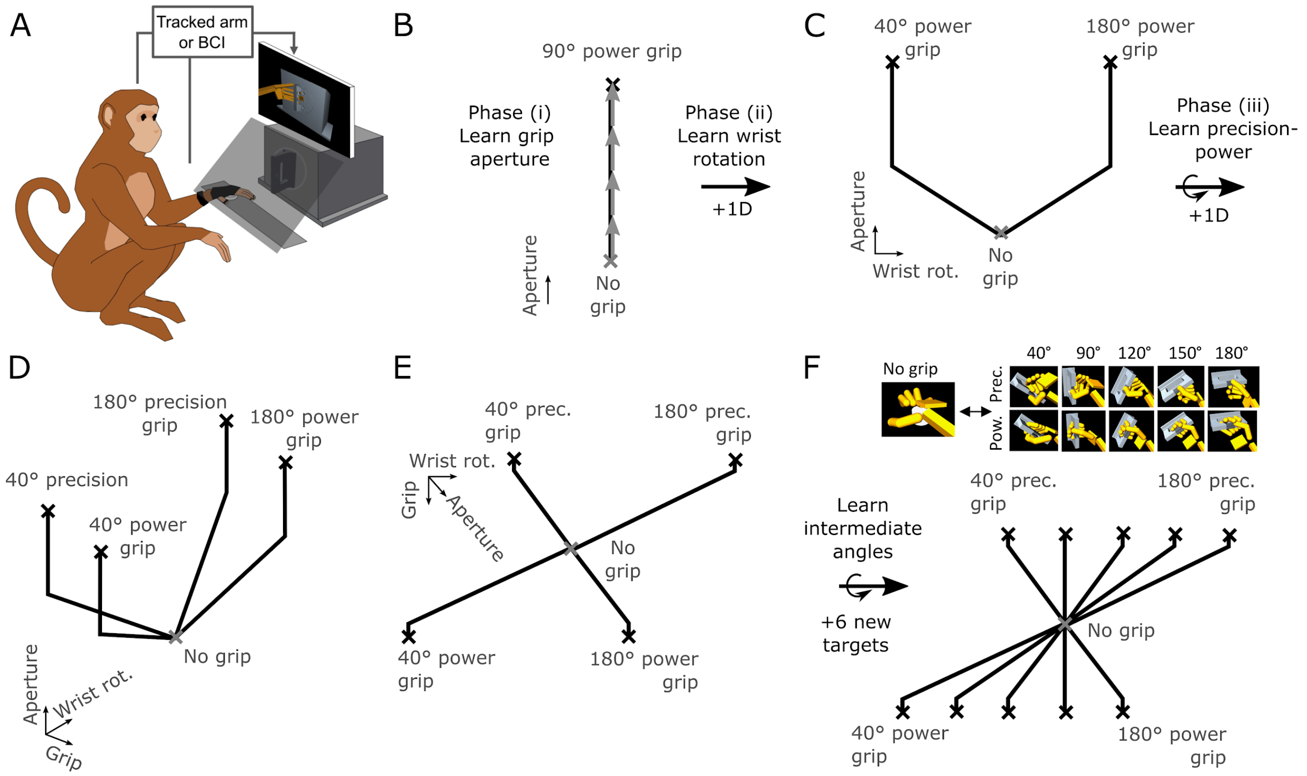

**Figure S1. Overview of the setup and trained degrees of freedom**

(A) Experiment setup. A subject sat in front of a screen displaying a virtual hand and a virtual rotating handle (implemented in MuJoCo). The virtual hand was controlled by either the native hand or the BCI. Neural data extracted from floating microelectrodes was used to control the BCI. During BCI control, lifting the hand to grasp would abort the trial. The native hand was concealed through a dark plate.

(B-F) Schematic of the training phases, degrees-of-freedom (DOF), and trajectories used to train the BCI. Note that the subject does not see these trajectories but only receives feedback from the grips on the screen.

(B) In Phase (i), subjects were trained to control only the grip aperture DOF by changing the hand posture from open to hand closed when reaching the 90° power grip. Full brain control was achieved in a few sessions.

(C) In Phase (ii), subjects learned to control wrist rotation DOF by achieving one of the two wrist angles required for a power 40° and 180° grip. Bents in the trajectories aim to emulate the curvatures in the native arm kinematics (Fig. S2, Methods).

(D-E) In Phase (iii), subjects learned to control a grip type DOF which selected the power or precision hand shape in the virtual hand. Together, the DOFs form a 3D space seen here for clarity from a side (D) and the top (E). Trajectories have an offset to compensate for the differences in wrist rotation for the power and precision grip for a given handle orientation.

(F) In addition to the training phases, subjects learned the intermediate target angles at the end of the experiments. For this, we asked the subjects to also grasp the handle at the 90°, 120°, and 150° orientations with both grip types. These were correspondingly, intermediate trajectories in the 3D space. Pictures in (E) display the virtual reality equivalent of the grips.

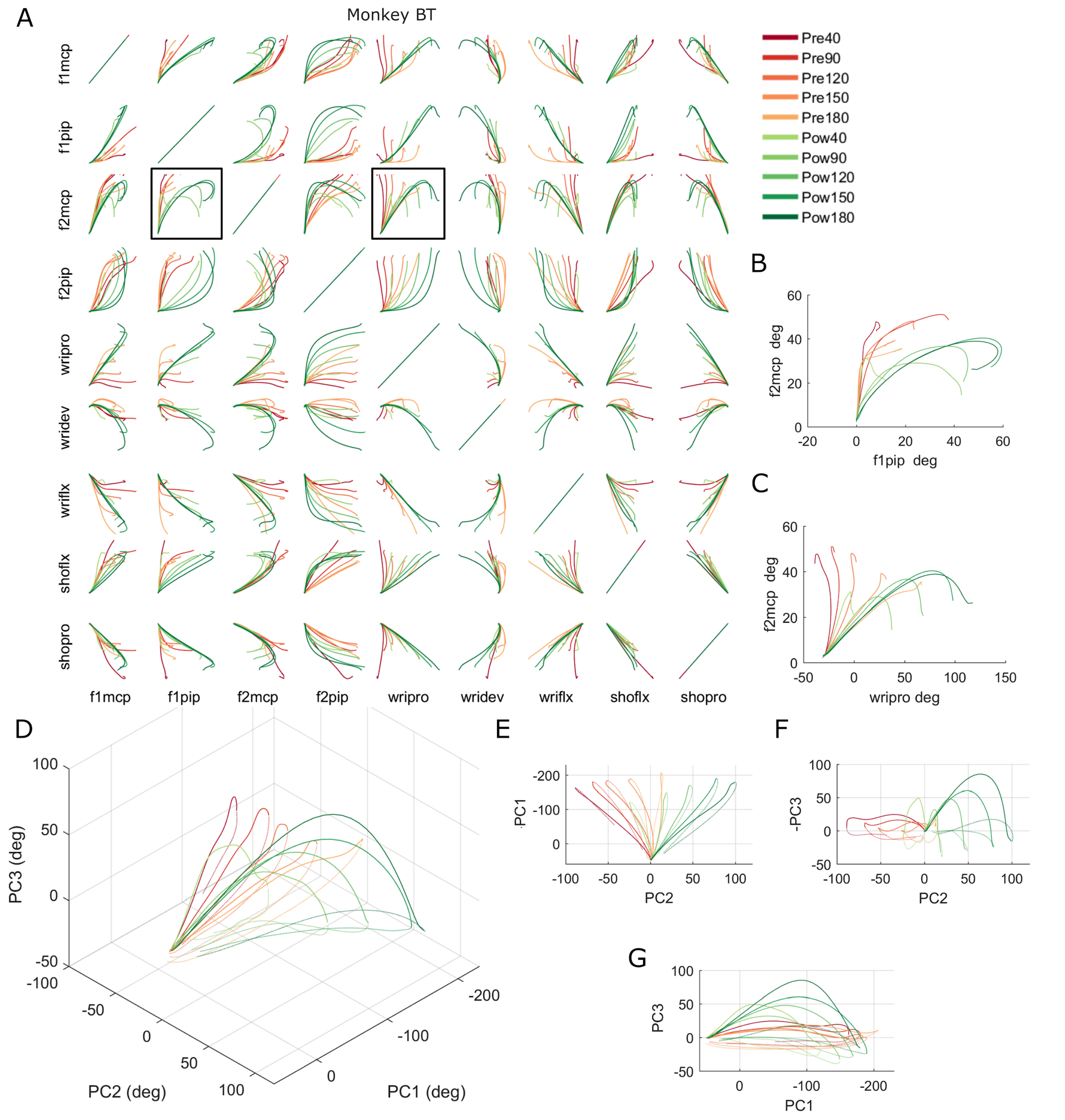

**Figure S2. Grip execution describes curved trajectories in the joint state-space**

(A) Example projections of the trajectories in joint angle space during native hand precision and power grips. Each sub-panel compares the evolution of a joint angle (out of 32 of the arm) from the neutral hand rest position (where the 10 traces converge) to the handle hold period (divergent traces end). During the execution of the grasp, joint angle pairs tend to describe curved trajectories (reflecting time differences in the evolution of each angle). Curvatures can be observed comparing arm and hand joint angle values or finger joint pairs. To understand these curvatures, imagine the combination of functions x=cos(t) and y=sin(t), with time parameter t. In 2D (x vs. y), the phase differences correspond to a curve (1/4 of a circle).

(B-C) Zoom in on the example pairs highlighted in dark boxes in (A). Axes are in degrees. (A-C) Pow: power grips, Pre: precision grips, f1: thumb, f2: index, mcp: metacarpophalangeal joint, pip: proximal interphalangeal joint, wri: wrist, sho: shoulder, flx: flexion, pro: pronation.

(D) Three first principal components from all joint angles (32 angles, >90% of the total variance). Solid line is the trajectory from hand rest to hold and dotted line is the return to rest phase of the movement. In general, movement in the kinematic space can be described as a curved trajectory. At hand rest, monkeys were usually in a neutral, relaxed position. During displacement to the handle (grip preparation) the whole hand was usually extended in preparation for the grip, and then the grip was executed. Upon return, monkeys usually "dragged" back the hand in a relaxed manner. The lesser curvature during the return phase indicates that most of the curvature is due to hand preparation which is not done when the animal returns the hand to rest.

(E-G) Upper and side views of (D). Data are trial averages from one entire recording session from monkey BT executing two grip types and all handle orientations.

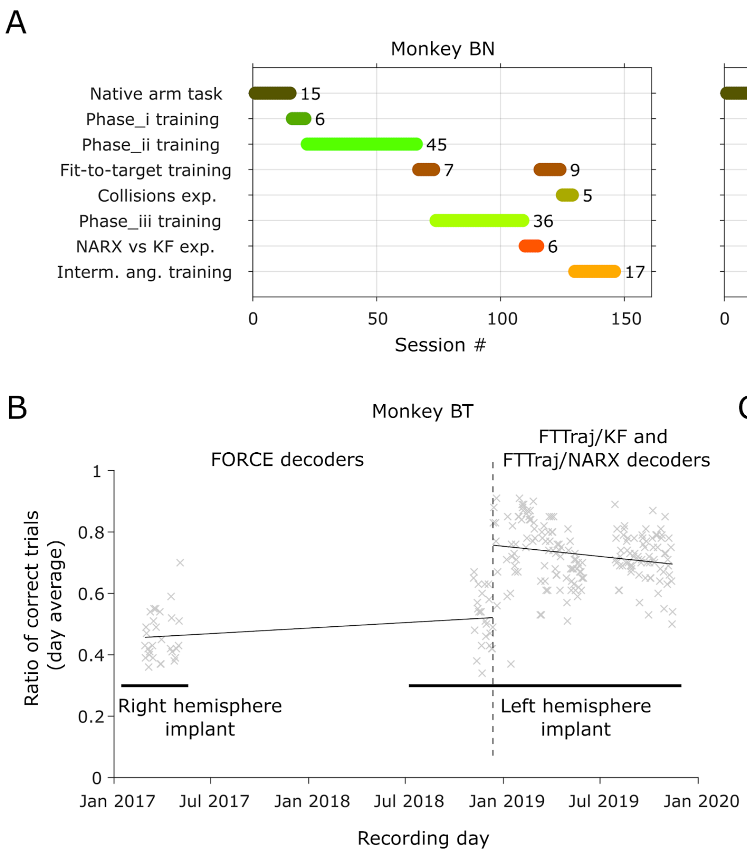

**Figure S3. Training timelines**

(A) Timeline of experiments. Rows represent a type of experiment and bars the number of sessions for each specific experiment or training period. Numbers next to each bar show the total session count for a given block. Sessions correspond to training or recording days. The graph also represents the order in which each phase and experiment was performed over time. The fit-to-target training of monkey BN was suspended after 7 sessions and later reinitialized as we decided the subject required more experience with the BCI to continue with the experiment. Note that training of the native grasping task is not shown here (it usually requires 3 to 6 months of training).

(B) Performance for all BCI recording sessions of monkey BT. Data includes the performance before starting training of our strategy (FORCE decoders) and after starting training our strategy (fit-to-trajectory position and velocity decoders). Before Jan. 2019, velocity-based decoders (FORCE RNN with 90% velocity and 10% position control) were used. Other FORCE configurations (e.g. 50% velocity and 50% position) made the decoder unstable and further worsened performance. For this reason and for training consistency, we always used the decoder fixed in the reported 90%-10% configuration. This performance difference seemed to be independent of the number of sessions and array quality, showing similar results in a ~4y old implantation and a recent implantation. A performance difference was immediately noticeable with decoder change. We did not attempt using the FORCE decoder with monkey BN.

(C) Example sessions comparing 10 decoder sessions of Monkey BN and 15 of Monkey BT using the fit-to-target KF with and without the "causal intervention" (CI, innovation 2 of Gilja et al.[1]). CI sessions were performed after the end of Phase (ii) training for Monkey BN and after the end of intermediate angles training for Monkey BT. Note that these sessions are not considered part of the main experiments and thus are not represented in (A). Bar represent the mean. p-value: Wilcoxon signed-rank test.

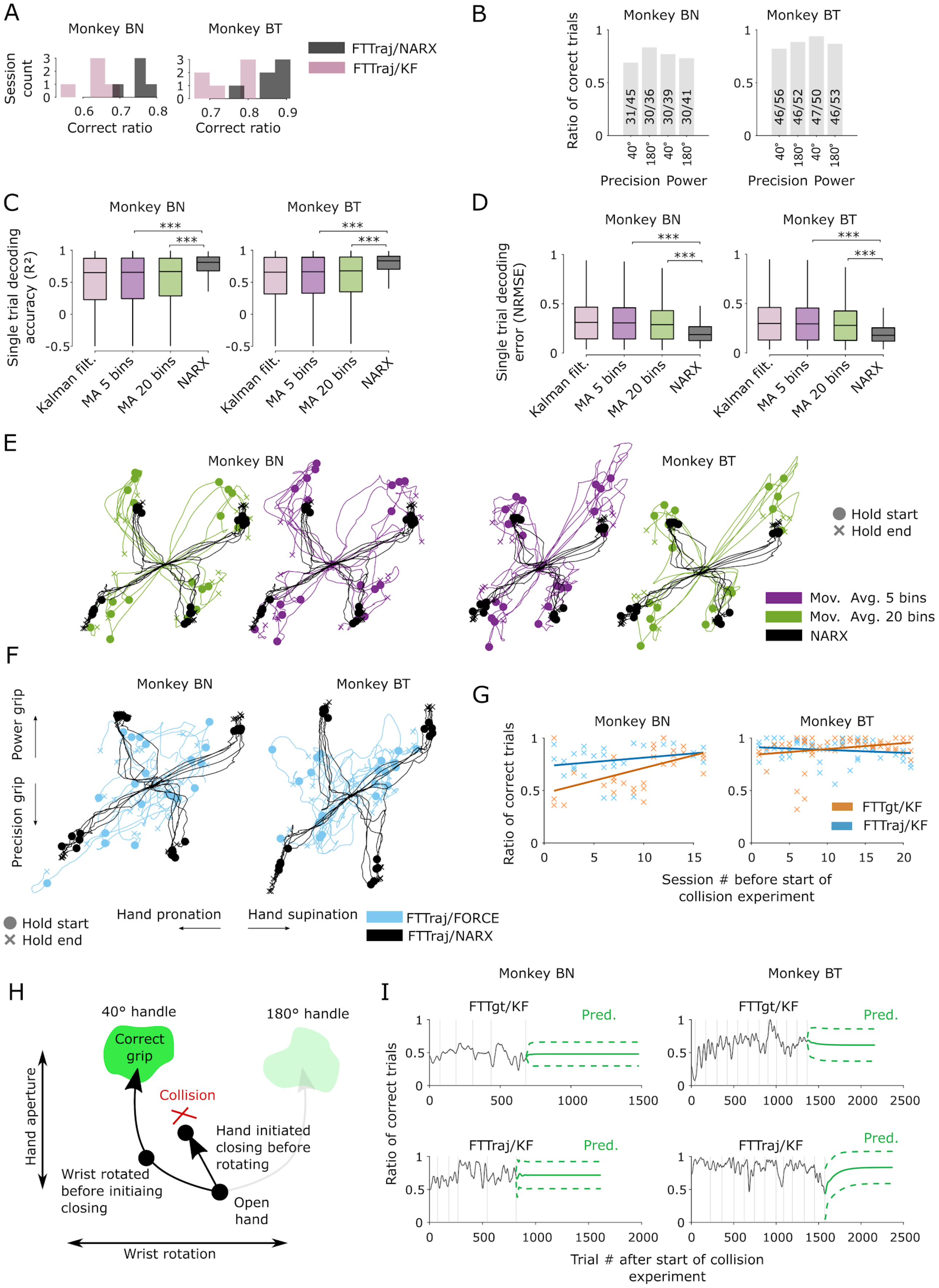
A
Monkey BN
Monkey BT
Session count
Correct ratio
FTTraj/NARX
FTTraj/KF
B
Monkey BN
Monkey BT
Ratio of corect trials
31/45
30/36
30/39
30/41
46/56
46/52
47/50
46/53
40°
180°
Precision Power
C
Single trial decoding accuracy (R²)
***
Kalman filt.
MA 5 bins
MA 20 bins
NARX
D
Single trial decoding error (NRMSE)
E
Hold start
Hold end
Mov. Avg. 5 bins
Mov. Avg. 20 bins
NARX
F
Power grip
Precision grip
Hand pronation
Hand supination
FTTraj/FORCE
FTTraj/NARX
G
Ratio of correct trials
FTTgt/KF
FTTraj/KF
Session # before start of collision experiment
H
40° handle
180° handle
Correct grip
Collision
Hand initiated closing before rotating
Wrist rotated before initiaing closing
Open hand
Hand aperture
Wrist rotation
I
FTTgt/KF
FTTraj/KF
Pred.
Trial # after start of collision experiment

**Figure S4. Performance of the fitting strategy**

(A) Distribution of individual session performances comparing our full strategy to the KF-Only variant. Same data as in Fig. 2.

(B) Performance per condition for one example session of each monkey during FTTraj/NARX control.

(C-E) Comparison of the NARX layer to plain smoothing.

(C,D) Single trial accuracy and root mean squared error of the decoded trajectory after replacing the NARX layer with a moving average filter (MA) of length 5 or 20. KF and NARX outputs are provided as a reference (***$p < 0.001$ Wilcoxon rank-sum test, Same data and N as in Fig. 2). Boxes: median, 25th and 75th percentiles, and range of the distribution.

(E) Example trajectories for the comparison of NARX to the MA filter. Same format as in Fig. 2.

(F) Example offline trials using the FORCE RNN (FTTraj/FORCE) compared to online trials using fit-to-trajectory with NARX (FTTraj/NARX). Data was produced offline using the neural signal during FTTraj/NARX control. In the FTTraj/FORCE case, hold start and hold end represent the periods where the corresponding FTTraj/NARX trial hold periods started and ended.

(G-I) Comparison of fit-to-target and fit-to-trajectory strategies.

(G) Performance during training previous to the collision experiment. Before the collision task, monkeys were trained to use the BCI with fit-to-trajectory and fit-to-target with limited collisions until they achieved similar performance. Before the start of the collision task, subjects had not been exposed to full collisions in the virtual environment. Every x represents a decoder run.

(H) Schematic of the collision task as seen in the wrist rotation and hand aperture DOFs. To properly achieve the 40 and 180° configuration the subject should coordinate the hand opening and wrist rotation such that the hand is opened enough when achieving the correct wrist angle (curved trajectory). Failure to prepare quickly enough would result in the hand being closed too early and a collision of the fingers with the handle (straight trajectory).

(I) Forecast of performance for the combined trials of 5 sessions of Monkey BN and 12 extended sessions (+5 from Fig. 2H) of Monkey BT during fit-to-target (top) and fit-to-trajectory (bottom) (40 steps moving average). Vertical gray lines show the session limits, green solid lines shows the prediction, and green dashed lines the 95% confidence intervals of an ARMA(4,4) forecast.

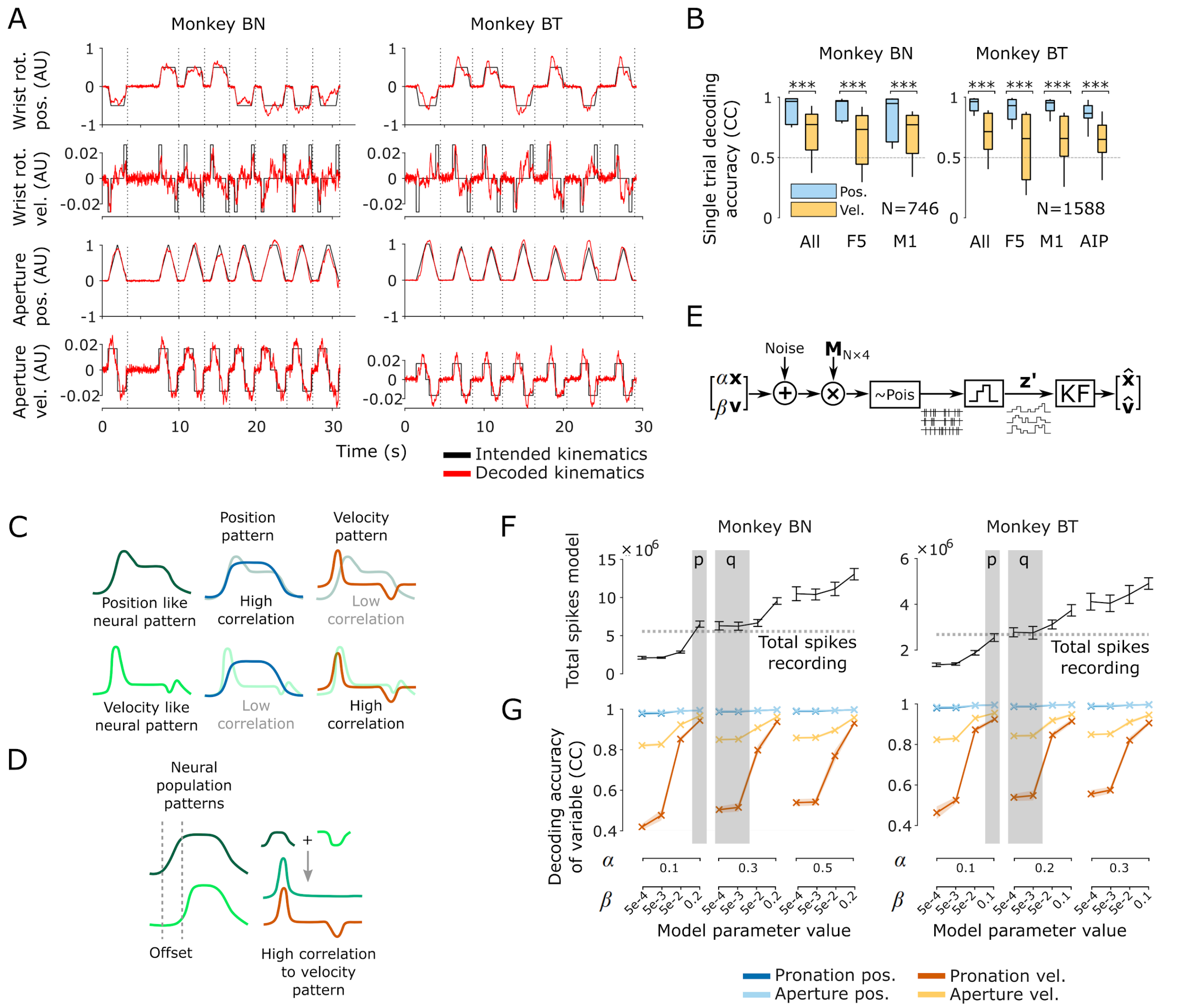

**Figure S5. Neural activity correlation to position and velocity during BCI control**

(A) Example decoded kinematics during BCI control. Data was taken from one representative session of Fig. 3A.

(B) Single trial decoding accuracy during BCI control of all areas combined and separated (same sessions as in Fig. 3A). We obtained CCs by comparing each decoded kinematic to the intended kinematic on each trial. Box plot: median, 25th and 75th percentiles, and distribution range. ***p<0.01, Wilcoxon signed-rank test on the single trial distribution.

(C,D) Schematic of possible unit-to-kinematic correlations.

(C) Neural patterns can resemble kinematic position patterns, or velocity patterns.

(D) However, velocity-like signals can also be produced in the population through linear combinations of position-like patterns with an offset. The dark green plus the inverted (negative) light green magenta produce a velocity-like pattern.

(E) Encoding model. Z-scored position (x) and velocity (v) signals are tuned with parameters α and β. The model injects noise and then maps the signals to a high-dimensional space of N=130 simulated units. The artificial neural signal was then Poisson discretized and binned (z'). A decoder (Kalman filter) was used to recover the original signal.

(F,G) Simulation results for different α and β. p, q denote two regimes in the parameter space where the simulated total spikes match those of the recordings. p: high-β regime. q: low-β regime.

(F) Average total spike count for each simulation. Dashed line: total spike count of the original BCI recording. Error bars: 2×STD.

(G) Correlation of decoded single-trials to trained target trajectories for different values of α and β using the encoding model on one BCI recording of monkey BN. For each point in the plot, we performed 10 simulations and plotted the average. Shades: 2×STD.

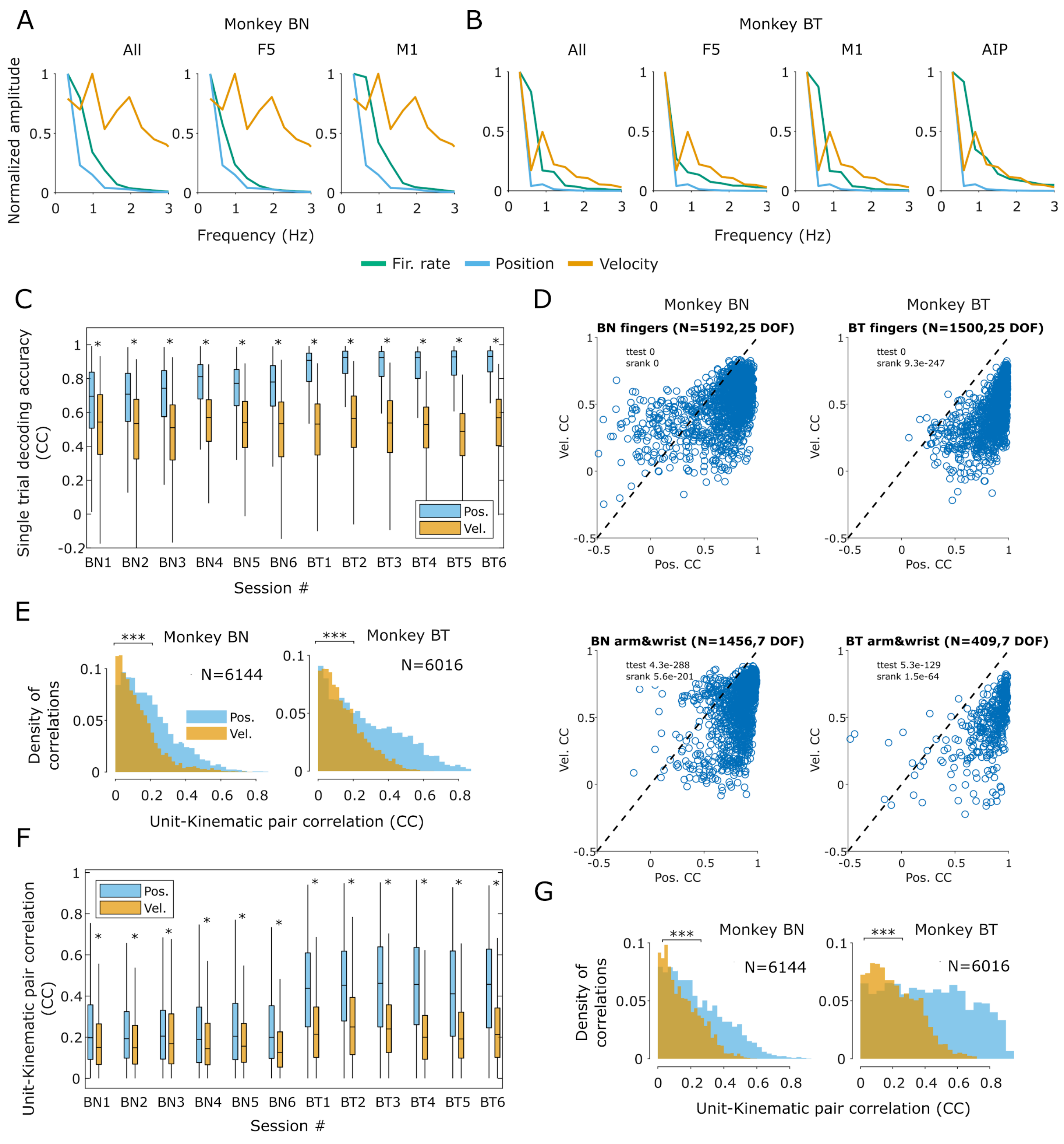

**Figure S6. Neural activity correlation to position and velocity during native grasp**

(A,B) Average of the Fourier transforms from all PETHs of one native arm recording of each monkey. Here, the frequency distribution of individual areas is compared to all areas.

(C) Position vs. velocity decoding accuracy on six native arm recordings of each monkey. Each position (32) and velocity (32) kinematic was predicted from neural activity on single trials using two independent KFs. $R^2$ was obtained by comparing to average kinematics. $R^2$ values below -0.5 were discarded. *p<0.001, t-test and Wilcoxon signed-rank test. Box plot: median, 25th and 75th percentiles, and distribution range.

(D) Same data as the native task single-trial decoding analysis of Fig. 3, but separating arm and hand joints. Top: fingers, 25 joint angles. Bottom: arm, 7 joint angles including the wrist. $R^2$ values below -0.5 were discarded.

(E) Pairwise correlations (abs. value) of all unit/position and unit/velocity PETH pairs for one native arm recording. Each data point in the distribution represents the CC of one unit PETH to one kinematic PETH. N: number of pairs. (***p<0.001, Wilcoxon signed-rank test).

(F) Same pairwise correlation analysis of Fig. 3, but now comparing six native arm sessions. PETHs were denoised here as in (I). CCs correspond to absolute values. *p<0.001, t-test and Wilcoxon signed-rank test. Box plot: median, 25th and 75th percentiles, and distribution range.

(G) Same data, colors, and pairwise correlation analysis as in (E), but only 90% of the variance was kept after PCA. N: number of unit/kinematic pairs. Distributions are significantly different (***p<0.001, Wilcoxon signed-rank test).

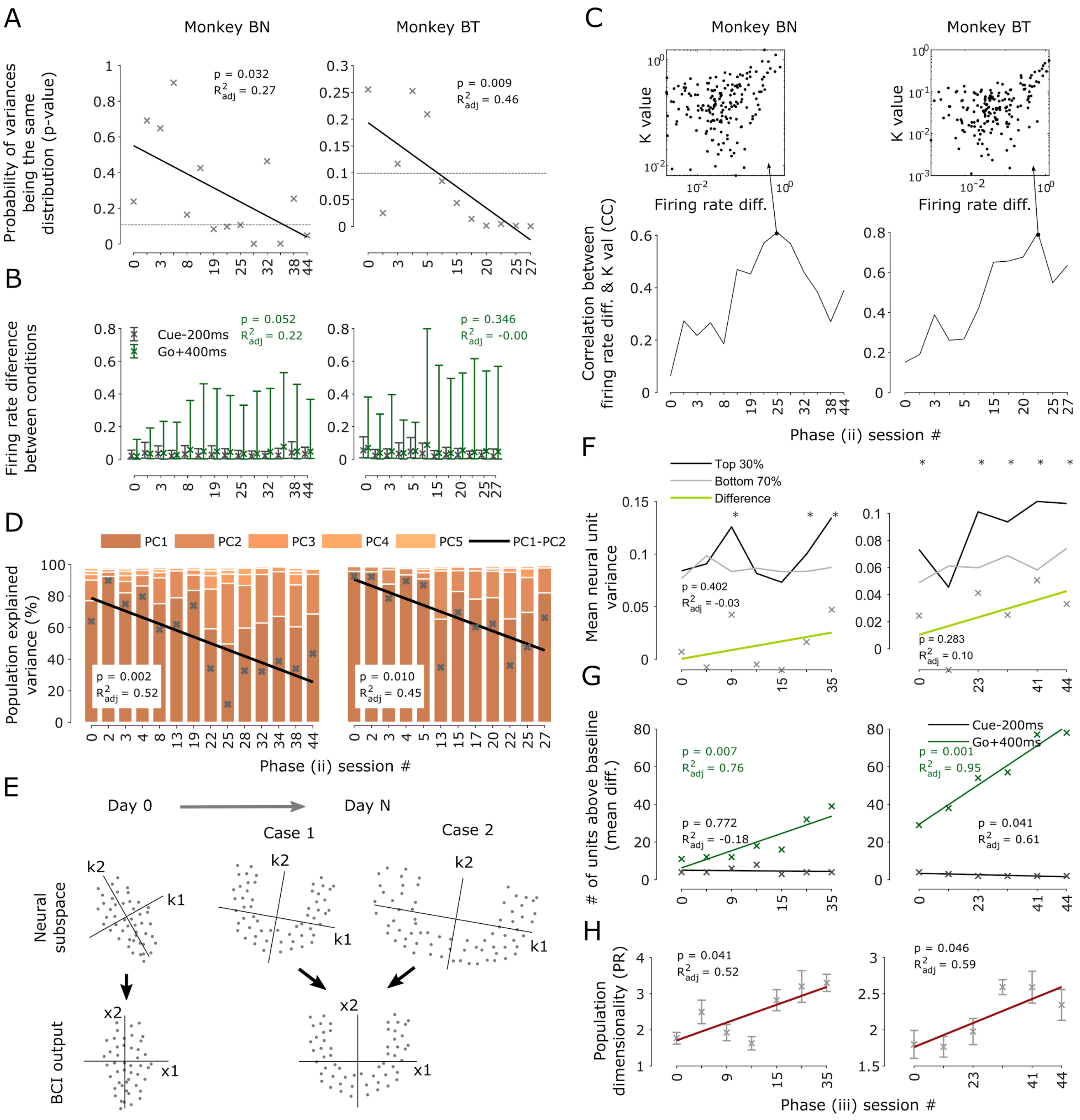

**Figure S7. Changes in population activity during training of Phase (ii) and Phase (iii)**

(A) p-values for the change in variance over sessions analysis of Fig. 4B. Dashed line indicates p=0.1.

(B) Approximate distribution of unit firing rate differences. The number of units of each session ranged between 130 and 180 (Methods). Rate differences were calculated from the condition-averaged firing rates (absolute values). We did not find a clear trend in the change of difference distributions over days. Stars: distribution means. Error bars: 10 and 90 percentile of the distribution.

(C) Over-session correlation between individual unit separation (whose approximate distribution is shown in (B)) and the weights the matrix K in the KF gives to each unit for the wrist rotation DOF (absolute values). Insets show the values used to obtain the correlation on the session indicated by the arrow. Here each point corresponds to a neuron.

(D) Redistribution of principal component explained variance with our adjusted PCA procedure. Change in the difference between PC1 and PC2 indicates that dimensionality of the neural population increased over sessions. x's represent the difference between PC1 and PC2 explained variance, and the regression line is a fit of the 'x' values.

(E) Motivation behind the scaling procedure used for dimensionality analysis. While PCA is sensitive to large variances, decoders are less sensitive. Neural population patterns (dots) are mapped to BCI outputs (x1,x2) through the row vectors (k1,k2) of the Kalman Matrix K. Here k1 and k2 map to position coordinates x1 and x2. Case 1: To produce kinematic separation, population activity might separate in a specific readout dimension (k1) proportional to other dimensions (k2). Case 2: However, it might also over-expand to compensate for noise variability breaking this proportion (variance along k1 is now proportionally larger to k2). Decoder readout is unaffected in Case 1. To properly measure dimensionality change in the population, we normalized population variance by the range of the kinematic readout (width of the kinematic workspace). Importantly, the scaling factor is kept constant for all sessions to prevent inflating the data as kinematics change over sessions.

(F-H) Same as analyses and format of Fig. 4 but for recordings of Phase (iii). Differences in this case were calculated from the power minus precision firing rate averages.

(A,B,F-H) Numbers indicate regression p-value and adjusted $R^2$. High p indicates that session # does not have a significant effect on the variable of interest.

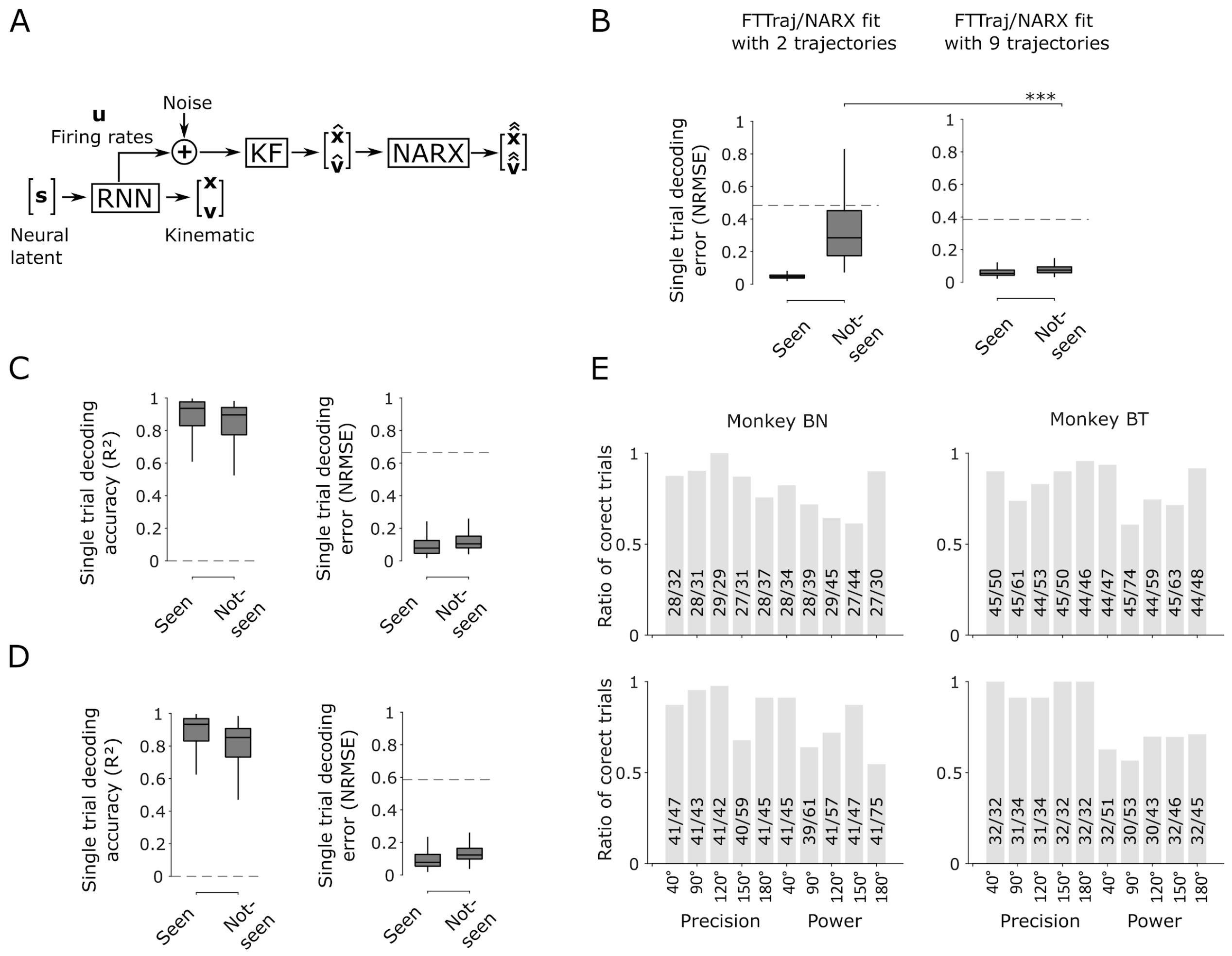

**Figure S8. Properties of the fitting strategy**

(A) Simulation model. Recurrent neural networks (RNN) were fit to produce kinematic trajectories (x, v) from neural latent trajectories (s). The simulated activations (u) were used to fit decoders using or our full strategy (x, v double hat).

(B) Effect of the number of trajectories used to fit the decoder on normalized root mean squared error (NRMSE). Left: FTTraj/NARX error if 2 trajectories are used. Right: FTTraj/NARX error if 9 trajectories are used. Seen: decoder error for trajectories used to fit the decoder (N=450 trials). Not-seen: decoder error for trajectories not previously seen by the decoder. (N=500 trials, ***p < 0.001 Wilcoxon rank-sum test).

(C) Decoder accuracy and normalized root mean squared error for a hypothetical 6 dimensional workspace (N=1350 seen and N=500 not-seen trials).

(D) Decoder accuracy and normalized root mean squared error for a hypothetical 12 dimensional workspace (N=1350 seen and N=500 not-seen trials).

(C,D) Dashed lines: chance levels (Methods). Boxes: median, 25th and 75th percentiles, and range of the distribution.

(F) Performance for four example sessions after monkeys learned the intermediate angles task. Bars: correct trials/total trials for a given condition.

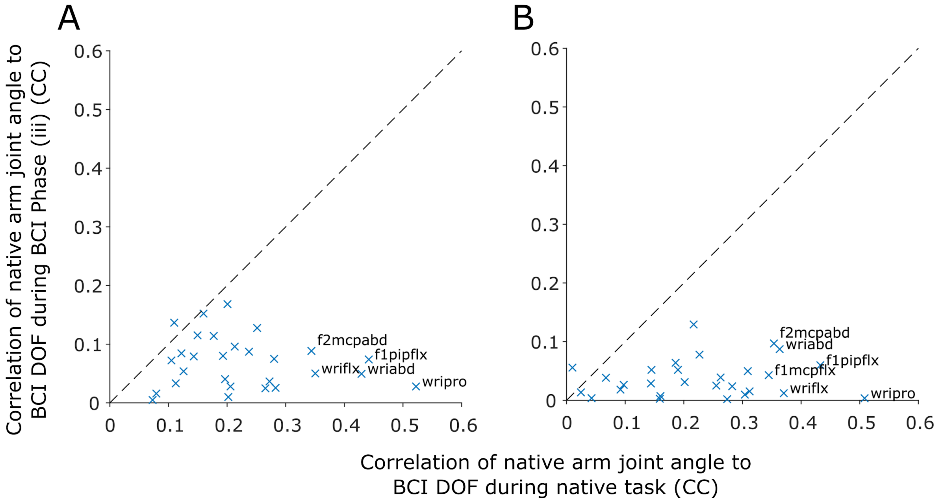

**Figure S9. Correlation of native arm joint angles to BCI output variables during two BCI experiments of Monkey BT**

(A,B) Two example sessions where monkey BT wore the glove while performing the BCI task compared to two native hand task sessions. Each x shows the average correlation between each of the 28 native hand angles during native hand use and the 3 BCI DOF outputs during brain control (trajectories in S1). To generate the x-axis, we produced artificial time and contion matched BCI trajectories and correlated them to joint angles during native hand control trials. To generate the y-axis we used the tracked small native hand movements during the BCI task and correlated them to the corresponding BCI output during BCI trials. Names highlight the joint angles with the highest x-axis correlation (same abbreviations as in Fig. S2).

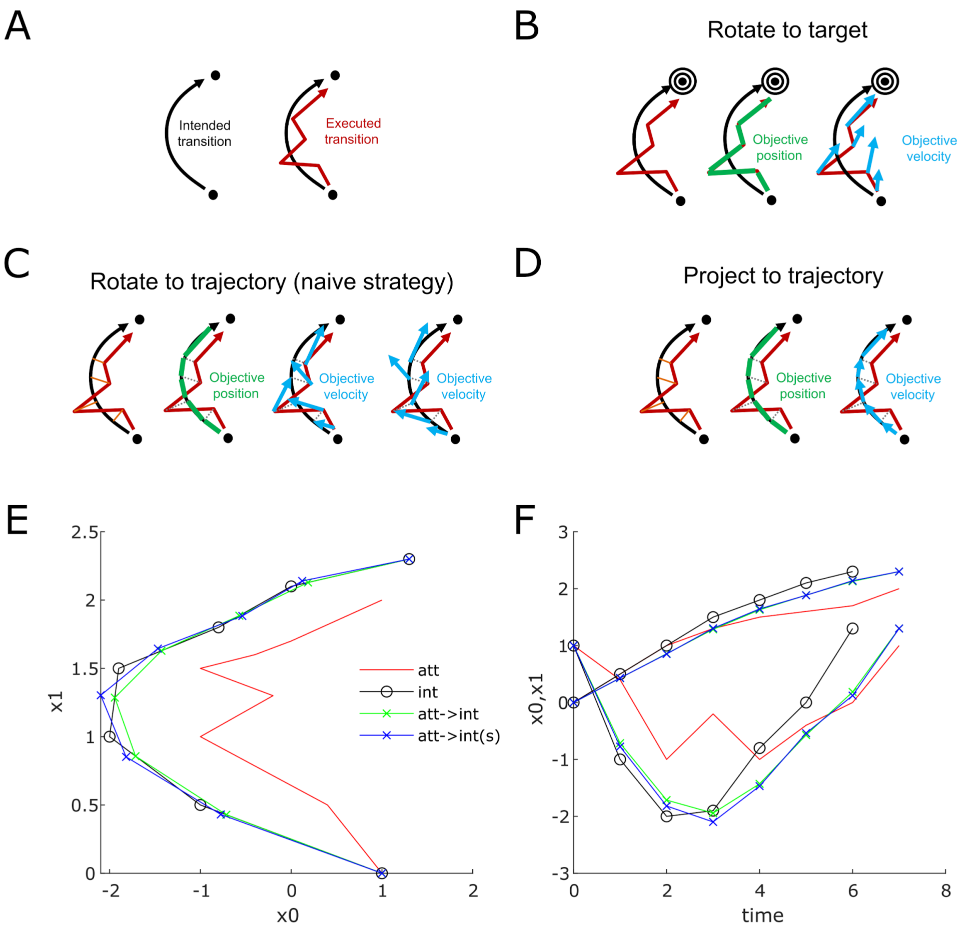

**Figure S10. Potential intention-estimation strategies and example implementation of fit-to-trajectory**

(A) We assume there is an intended trajectory (ideal hand configuration transition) and an executed trajectory (obtained from data from a decoded initial attempt) in some kinematic space. To recalibrate the BCI, a new objective position and a new objective velocity are needed as input for the decoder fitting procedure.

(B) The ReFIT strategy[1] assumes that the intention is to reach a specific target in the kinematic space. It is assumed that the intent of the subject was to move towards the target and the velocity vectors are redirected to the target. In ReFIT, the objective position is left unchanged as this does not influence the decoder (second innovation).

(C-D) When trajectories are relevant, we opted for projecting the attempted trajectory onto the intended trajectory to preserve the position intent and the velocity intent. The objective position is clear in this case as this will be along the intended trajectory. We determined two possible strategies to reorient the velocity vectors: (C) towards the trajectory, or (D) along the trajectory. Rotating towards the trajectory implies that the intended velocities shoot out of the trajectory when position is considered (C, rightmost diagram). We opted for strategy (D).

(E) Example fit-to-trajectory procedure for a two dimensional workspace (x0 vs x1). To project the trajectory preserving the shape of the intended trajectory, an interpolation method can be used (e.g. linear or spline). In our case, we used a linear method. Legend: att: attempted trajectory, int: intended trajectory, att->int: attempted onto intended trajectory (using linear interpolation, see example implementation), att->int(s): attempted onto intended trajectory (using spline interpolation, see example implementation).

(F) Example kinematic variables x0 and x1 over time. Note that this strategy preserves the timing of the attempted trajectory.

#### Supplemental tables

```
clearvars;
% Example trajectories
patt = [1 0; 0.4 0.5; -1 1; -0.2 1.3; -1 1.5; -0.4 1.6; 0 1.7; 1 2];
pint = [1 0; -1 0.5; -2 1; -1.9 1.5; -0.8 1.8; 0 2.1; 1.3 2.3];
Natt = size(patt,1)-1;
Nint = size(pint,1)-1;

% Prepare parameters
Laoi = Nint/Natt;
Tatt = 0:Natt;
Tint = 0:Nint;
Taoi = 0:Laoi:round(Natt*Laoi);

% Find parameters on the curve
paoi = interp1(Tint, pint, Taoi);
paois = interp1(Tint, pint, Taoi, 'spline');

% Plot
figure(1); clf;
subplot(1,2,1); hold on;
plot(patt(:,1),patt(:,2),'r'); plot(pint(:,1),pint(:,2),'-ok');
plot(paoi(:,1),paoi(:,2),'-xg'); plot(paois(:,1),paois(:,2),'-xb');
xlabel('x0'); ylabel('x1');
legend({'att','int','att->int','att->int(s)'},'Location','e');
subplot(1,2,2); hold on;
plot(Tatt,patt,'r'); plot(Tint,pint,'-ok');
plot(Tatt,paoi,'-xg'); plot(Tatt,paois,'-xb');
xlabel('time'); ylabel('x0,x1');
```

**Table S1. Example implementation of the fit-to-trajectory method**

The output of this Matlab code can be seen in Fig. S10.

| Comparison<br>(A vs. B) | Experience using A before test | Experience using B before test | Performance | Notes |
|---|---|---|---|---|
| Native arm vs. KF | Years for both subjects | 94 sessions (BN) (Fig. S2A)<br>110 sessions (BT) (Fig. S2A) | A > B<br>(Fig. 2B-F) | |
| Native arm vs. KF+NARX | Years for both subjects | 94 sessions (BN) (Fig. S2A)<br>110 sessions (BT) (Fig. S2A) | A > B<br>(Fig. 2B-F) | |
| KF vs. KF+NARX | 94 sessions (BN) (Fig. S2A)<br>110 sessions (BT) (Fig. S2A) | 0 sessions for both subjects | B > A<br>(Fig. 2B-F) | B was at a disadvantage but still was superior. |
| FORCE vs. FTTraj/KF | >50 sessions (BT) | 0 sessions (BT) | B > A<br>(Fig. S3B) | B was at a disadvantage but still was superior from the start and over sessions. |
| Fit-to-target vs. Fit-to-trajectory | 13 sessions (BN)<br>21 sessions (BT) | 14 sessions (BN)<br>20 sessions (BT) | B > A<br>(Fig. 2H-I, Fig. S4G-I) | Similar sessions and performance matched before test of collisions. |

**Table S2. Summary of native arm and online decoder comparisons**

### Supplemental notes

#### Collision experiment

Trying to understand the advantages of our fit-to-trajectory approach over previous approaches (Gilja et al.[1]), we designed a low-collision/full-collision task which required the subject to acquire the grasp target while preventing collisions with the handle. We trained the BCI in a low-collisions virtual environment where only the fingertips and the precision button were collidable (i.e. handle, finger and dorsum were not collidable by default, Fig. 2G). After completing Phase (ii) of the training, we further trained both subjects to perform the task using fit-to-trajectory (FTTraj) or fit-to-target (FTTarg) until achieving similar performance in both of these settings (Fig. S4E). During FTTarg, we used the grip configuration at hold as the end-target, following Gilja et al. Importantly, we used only the first ReFIT innovation, as the second innovation caused a performance drop in our case (Fig. S3C). After this, we introduced full collision in the virtual environment (i.e., handle, rest of fingers, and hand dorsum collidable, MuJoCo engine) at the same time for both settings. The environment required the subject to prepare for grasping by opening the hand on time to prevent colliding with the handle (Fig. S4H). Without prior experience with the collision environment, subjects were more successful using the fit-to-trajectory strategy (Fig. 2H,I). We did not observe any improvement in task performance while using the fit-to-target approach over the tested number of sessions (Fig. S4I).

There are caveats to this result. A direct comparison to full ReFIT (i.e the two innovations on Gilja et al.) was not possible in our case for several reasons:

1) From the start of training, we observed a lack of velocity control, demonstrated by the low performance of the FORCE decoder (Fig. S3B) and likely, by the performance drop seen with the second ReFIT innovation (Fig. S3C). For this reason the second innovation was disabled, and our comparison was not a full ReFIT vs. our method, but fit-to-target vs. our method as the figure labels state.

2) It is harder to establish clear end and intermediate target points when controlling a multi joint effector. While in a 2D cursor workspace, end targets are clear points and intermediate targets have a clear 2D designation (e.g. empty spaces next to an obstacle), in a multi-dimensional hand joint state-space, targets and intermediate targets are visually less distinguishable (e.g., configuring the hand to grasp). In our opinion, this makes the concept of ”targets” not very useful for tasks like ours.

3) The intricacies of monkey and BCI training allowed us to train only a single protocol from the start. We tried to compensate for these deficiencies by training subjects for several sessions to use fit-to-target (Fig. S4G). A more exhaustive comparison would have implied further training subjects to use fit-to-target in the collision task or different monkeys. Our results therefore only demonstrate that fit-to-trajectory was advantageous at the beginning of the collision task.

Our main point with this comparison is that the concept of “rotating to a target” does not extend to cases where the task objective is to produce kinematic trajectories, which is a requirement for any type of position based control. On these grounds, the comparison of full ReFIT to our method could be considered an “apple to oranges” comparison. ReFIT is likely very useful when the task goal is to control velocity, but differences in information content in different brain areas will likely impose different fit type requirements.

#### Area differences

Frequency analysis of the native task neural data by brain area indicated that area F5 had a lower amount of high-frequency components compared to M1 (Fig. S6A,B), and these frequency content differences seem to predict decoding quality for velocity of each area (Fig. 3G). Low high-frequency content in neural activity matches with lower velocity decoding quality in area F5 for monkey BT, and higher high-frequency content in neural activity matches with improved decoding quality in area F5 for monkey BN. Higher high-frequency content in area M1 corresponds to improved velocity decoding quality in area M1 for both subjects. These observations match the role of these two areas in the grasping circuit[2], F5 activity being closer to grasp types and thus to posture information, and M1 being closer to the output of the circuit and thus containing both low- and high-frequency commands required for muscle control. We have been able to decode high-quality kinematics from AIP in past work[3], but the availability of high-frequency information in AIP requires further study.

##### Restrictions to hand movement during BCI control

We trained our subjects to perform the BCI task without lifting or grasping with the native hand. Simultaneous movement of the hand is not detrimental to BCI use and has been used in previous protocols[1,4] (although it can cause interference depending on the decoder type[5]). During BCI use, subjects performed small contractions of the hand. Although we did not measure EMG, we believe these contractions had limited influence on the task. These movements had a very low correlation with the BCI output (Fig. S9, Methods). If stereotypical movements had control over the task (e.g. moving the hand slightly to the left or right), it would have been faster for the monkeys to learn the 2D task. In our task, we opted for preventing lifting of the hand as we were generally interested in the plasticity changes required to control this ”new” effector. The neural changes measured in our data seem to indicate that control of the task required a neural adaptation to correctly separate conditions. The time it took to learn the task indicates this adaptation could have been a skill learning process[6,7].
